# Spatiotemporal Dynamics and Migration Pathways of the Transboundary Beet Webworm *Loxostege sticticalis* (Linnaeus) Between Northern China and Mongolia in 2022

**DOI:** 10.64898/2025.12.28.696789

**Authors:** Xing-Yue Pu, Yi-Yang Zhang, Hai-Bin Gu, Rui Zhong, Gui-Jun Wan, Fa-Jun Chen, Qiu-Lin Wu

## Abstract

Clarifying the population dynamics, source-landing relationships, and migration pathways of the beet webworm *Loxostege sticticalis* (Linnaeus) both domestically and internationally, as well as understanding the meteorological mechanisms shaping these patterns, is pivotal for remote, accurate, and location-specific pest forecasting. Based on light trap data from northern China and field survey data from Mongolia in 2022, we simulated the migration trajectories, source regions, and primary landing areas of *L. sticticalis*, and analyzed the synoptic systems and processes during its migration. The results indicate frequent exchange of *L. sticticalis* populations between China and Mongolia in 2022. The *L. sticticalis* migrants initiating their flights from Mongolia primarily undertook a southeastward migration pathway, supplemented by eastward ‘cyclonic’ and southwestward paths. The main landing areas were located in North China and Northeast China, with migration events potentially extending to Shandong, Heilongjiang, and Xinjiang provinces. Populations originating from North China exhibited a capacity for migrating into Northeast China and Mongolia through 1 to 5 consecutive nights of flight. During this period, the Northeast China Cold Vortex (NCCV) and the Mongolian Cyclone alternately regulated the synoptic circulation pattern governing the migration of *L. sticticalis*. The spatiotemporal distributions and intensities of these systems were key determinants of the transboundary migration routes and distances of *L. sticticalis*. The NCCV dominated, and the precipitation and downdrafts it induced were crucial for the massive landing of *L. sticticalis* in northern China.

## 1. Introduction

The beet webworm *Loxostege sticticalis* (Linnaeus) belongs to the Pyralidae family in the order Lepidoptera. It is characterized by its migratory ability, destructive nature, periodic outbreaks, and omnivorous diet [1], with its range mainly spanning the narrow strip of Eurasia between 36° N and 55° N, including countries such as China and Mongolia [2,3]. *L. sticticalis* infests more than 50 host plant species across 21 families, mainly damaging crops such as soybean and sunflower, and poses a serious threat to agriculture and animal husbandry in China’s Three North Regions [4]. During severe outbreak years, it can inflict yield losses ranging from approximately 60% to a staggering 100% [5,6]. *L. sticticalis* ranks third among Category 1 crop pests, as announced by the Ministry of Agriculture and Rural Affairs, following only *Spodoptera frugiperda* and *Locusta migratoria*.

China has been expanding planting acreage, boosting the planting of soybean and other host crops, and reducing the planting of maize as part of its national goals, such as the Rural Vitalization Strategic Plan. This may help *L. sticticalis* populations spread [7]. Since the founding of the People’s Republic of China, *L. sticticalis* has undergone three major outbreak cycles: 1955–1961, 1978–1984, and 1996–2009 [8]. After the third cycle, infestations remained at low levels from 2010 to 2016 [9]. In 2018, however, high densities of first-generation larvae were detected in the border regions of Inner Mongolia, Heilongjiang, and Jilin [10], followed by a marked increase in overwintering adults in 2019 [11]. The population regained strength in 2020, signaling the initiation of a new severe outbreak cycle [12]. Additionally, global warming has intensified the northward expansion of this pest’s overwintering range in North and Northeast China, and promoted the westward migration of its second- and third-generation adults to high-altitude areas [13]. The distribution of *L. sticticalis* adults in North China is widespread and severe, with the activity regions of pests from the overwintering generation to the first generation advancing northward [14]. In Northeast China, the peak trapping period for overwintering-generation adults has come earlier, with large moth populations, prolonged duration, and multiple migration peaks [15]. Contrary to the conventional view in the 1980s, Chen Xiao et al. [6] proposed that North China serves as a source of migrant *L. sticticalis* populations for Northeast China, but not the primary source. The pest populations primarily originate from local overwintering groups and those from neighboring countries such as Mongolia. Part of them come from the border areas between China and Mongolia, China and Russia, or the tri-border region of the three countries [16,17]. *L. sticticalis* has long been widely distributed in Russia, Mongolia, and other regions [18]. In 2008, although the population of first-generation larvae in China was small, a massive influx of first-generation adults migrating from Russia and Mongolia resulted in a severe outbreak of second-generation larvae [19]. Moreover, Mongolia shares a long border with China, and contains similar vegetation types which *L.sticticalis* primarily feeds [20].

Long-distance migration is initiated within favorable backdrop wind fields, while it can be seriously hindered by low-level winds, low-level jets, and, particularly, severe air convection at the migration altitude [21,22]. Cyclonic systems can significantly disrupt background wind fields. For instance, when a long-wave trough in the westerlies deepens to a certain extent, the air exchange between the cold trough and its cold air source is blocked, forming a low-pressure center south of the trough, known as a cutoff low [23]. The cutoff low in Northeast China, designated as the Northeast China Cold Vortex (NCCV), is a persistent and relatively stable atmospheric system present between 35–60° N and 110–145° E, regularly occurring from May to August each year [24]. This spatial and temporal range coincides with the migration period and geographic distribution of *L. sticticalis*. Chen Xiao et al. [25] stated that the low-level winds impacting the migration direction of *L. sticticalis*, along with precipitation and severe convective weather leading to its mass landings, may derive from the NCCV. The NCCV, typically accompanied by squall lines, storms, and heavy rain [26,27], gives rise to prolonged rain, low temperatures, and abrupt severe convective weather in Northeast China and Inner Mongolia [28,29]. This pest is forced to land when the temperature falls below 15 °C or the cumulative precipitation across 12 h reaches 0.1 mm [30,31]. Low temperatures and rain can also have a significant effect on the growth and production of crops [32,33], thereby indirectly affecting how often outbreaks of *L. sticticalis* occur and how much harm they cause. Previous investigations on this pest have mainly been conducted within China, and the spatiotemporal connectivity between domestic and international migratory populations has yet to be fully revealed.

Therefore, based on the population dynamics of *L. sticticalis* in Mongolia, it is of crucial importance to identify the overseas origins of the pests in China by simulating its migration trajectories, source areas, and major landing regions, and analyzing the associated weather systems and processes during migration. In this research, we used the Hybrid Single-Particle Lagrangian Integrated Trajectory (HYSPLIT) model, a well-known insect migration trajectory model, in conjunction with ArcGIS 10.7, to predict the migration pathways of *L. sticticalis* populations in China and Mongolia in 2022. A statistical analysis of landing places was conducted to examine the connectedness of populations between the two nations. Additionally, Python software 3.10.18 was utilized to analyze the meteorological conditions influencing *L. sticticalis* migration to elucidate its migration processes and landing mechanisms, thereby providing a theoretical foundation for the prediction, early warning, and control of this pest.

## 2. Materials and Methods

### 2.1. Pest Occurrence Data

The pest occurrence data, including high-light trap records, conventional light trap records, and female moth ovary dissection results from northern China in late June 2022, as well as field survey data (number of moths per hundred steps) from Mongolia between June and August 2022, were provided by the Pest Forecasting Division of the National Agro-Tech Extension and Service Center. The monitoring data are shown in Figure 1.

**Figure 1.**
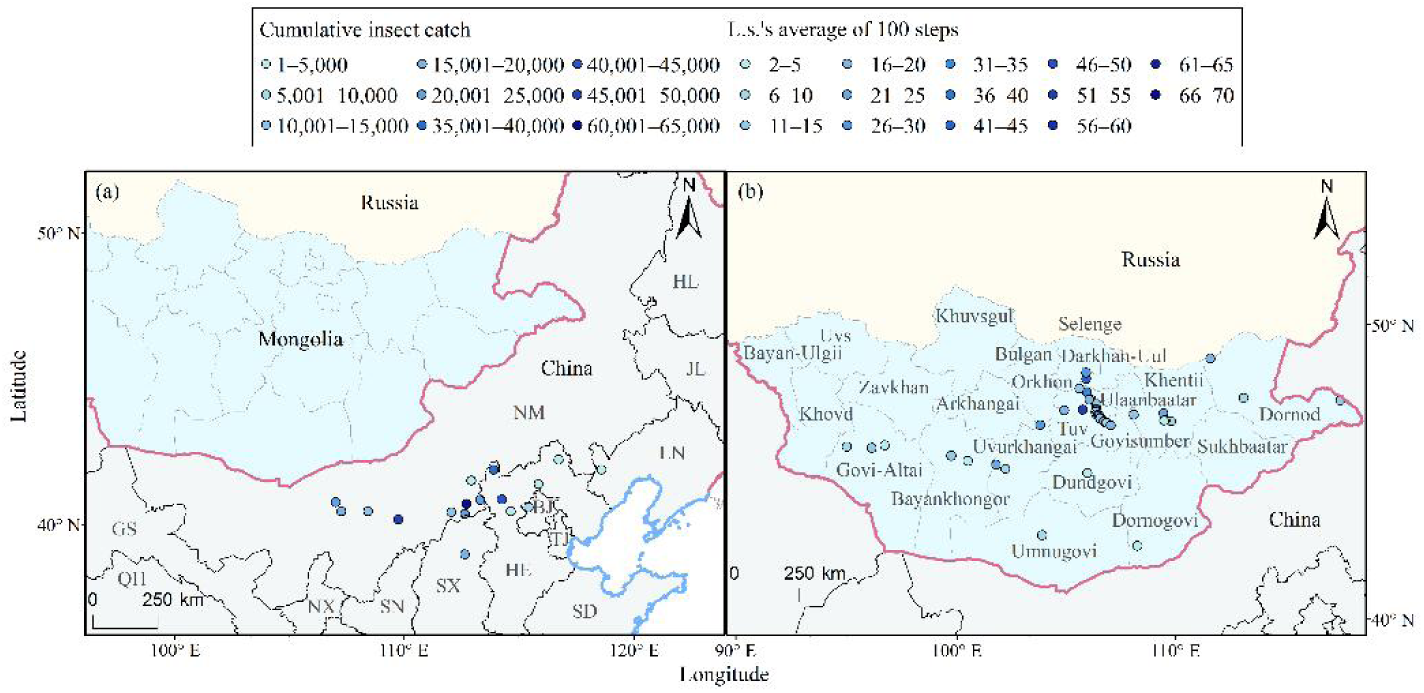
Monitoring of *L. sticticalis* in northern China (**a**) and Mongolia (**b**). In panel (**a**), provincial-level administrative divisions are labeled with initial abbreviations: HL—Heilongjiang Province; JL—Jilin Province; LN—Liaoning Province; NM—Inner Mongolia Autonomous Region; BJ—Beijing City; TJ—Tianjin City; HE—Hebei Province; SD—Shandong Province; SX—Shanxi Province; SN—Shaanxi Province; NX—Ningxia Hui Autonomous Region; GS—Gansu Province; QH—Qinghai Province.

### 2.2. Meteorological Data

The study utilized the ERA5 dataset (ECMWF Fifth Generation Reanalysis; https://cds.climate.copernicus.eu/ accessed on 16 December 2024), a global climate reanalysis product provided by the European Centre for Medium-Range Weather Forecasts (ECMWF), to extract meteorological data from 28 May to 13 August 2022, with a spatial resolution of 0.25° × 0.25° and a temporal resolution of 1 h. For this study, we selected the spatial domain of 85–140° E, 30–60° N—covering Northeast China, Northwest China, North China, and Mongolia—as the specific research area. Given that *L. sticticalis* migration typically occurs at night, we conducted meteorological analyses using Python, focusing on meteorological elements/conditions from 18:00 to 06:00 Beijing Time (UTC+8) on the following day. In the study area, the average elevations of monitoring stations are about 1000 m in China and 1335 m in Mongolia, with stations below 800 m accounting for less than 12.5%. We decided to examine predominant factors including the wind field (u and v components), vertical velocity, geopotential height, temperature at the 850 hPa level, and surface precipitation [1]. We utilized the geopotential height at 850 hPa to identify cyclones in Mongolia [34], while 500 hPa geopotential height and temperature were the criteria for defining the NCCV [24].

### 2.3. Simulating Migration Paths Using the HYSPLIT Model

In this work, we utilized the HYSPLIT model (https://www.ready.noaa.gov/HYSPLIT.php, accessed on 4 December 2024) to simulate the migration paths of both domestic and immigrant populations of *L. sticticalis*, thus identifying potential source areas and landing zones [35–37]. Cooperatively produced by the National Oceanic and Atmospheric Administration’s (NOAA) Air Resources Laboratory and the Australian Bureau of Meteorology, the HYSPLIT model has been used not only to trace the spread of air masses in order to understand their paths and other intricate processes including transit, transformation, diffusion, and deposition, but also to simulate the migration paths of different insect groups, such as *Aphidoidea*, *Nilaparvata lugens*, and Noctuidae [38–41].

Based on the sudden surge in *L. sticticalis* moth catches recorded by high-light traps or conventional light traps in China, we determined the characteristics of the populations according to the results of ovarian dissection in female moths. Specifically, populations peaking under light traps were classified as emigrants if the proportion of females with ovarian development at levels 3–4 was ≤65%, and as immigrants if otherwise [42,43]. Forward trajectories were used to identify the migratory pathways and main landing areas of *L. sticticalis*, while backward trajectories were employed to analyze the potential source regions of this pest. For stations lacking conditions for ovarian developmental stage analysis, populations were classified based on historical occurrence data of *L. sticticalis*. Additionally, existing research findings indicate that the preoviposition periods are 5.13 days for migratory populations and 8.70 days for non-migratory populations. It is important to note that individuals captured in the field may not have emerged on the same day; for instance, some adults captured on 18 June may have emerged around 14 June [44]. According to Frolov et al. [18], the overwintering generation of this pest in Mongolia begins to appear from mid- to late May. To analyze the migratory pathways of overseas populations, we defined the migration window as date(s) with reported moth count >1 individual per hundred steps and the 4 d before and 4 d after this date. Forward trajectories of emigration from each survey location, for a total of 9 d between 28 May and 13 August 2022, were simulated accordingly.

*Loxostege sticticalis* typically takes off around sunset, migrates during the night, and lands at dawn, with a flight duration of approximately 10 h, and migration altitudes between 300–500 m above ground level (AGL), with the core insect layer centered around 400 m AGL. Radar data indicate a scarcity of individuals over 700 m AGL, with moths typically flying for 1–5 nights [35,45]. In this study, we simulated the migratory routes of *L. sticticalis* based on specific biological traits and computational parameters: (1) Migration simulations were conducted for 1 to 5 consecutive nights, with 20:00 Beijing Time (UTC+8) serving as the starting point for forward calculation and endpoint for backward calculation, and vice versa for 06:00 the next day. (2) The endpoint of each night’s trajectory served as the starting point for the following night, and flight duration for each night could last a maximum of 10 h. (3) Flight altitudes were set at intervals of 100 m between 200 and 700 m. (4) During each night’s flight, it was assumed that *L. sticticalis* could potentially land at any full hour. To fully estimate the migratory process of *L. sticticalis*, we did not incorporate environmental factors such as host crop availability, high-altitude temperature dynamics, or hourly precipitation, which could actively or passively interrupt the migration process. Based on historical occurrence data of *L. sticticalis*, effective source regions in the backward trajectory results were further filtered (Table S1). Finally, all landing points were statistically analyzed and mapped using ArcGIS 10.7. The standard map of China used in this study (Map Review No.: GS (2024) 0650) was downloaded from the National Platform for Common Geospatial Information Services (https://www.tianditu.gov.cn/).

## 3. Results

### 3.1. Analysis of the Population Dynamics of Adult Loxostege sticticalis in Northern China

According to the monitoring data recorded by high-light traps and conventional light traps, overwintering-generation adults of *L. sticticalis* began to appear successively in regions such as Inner Mongolia, Hebei, and Beijing from late May 2022. The number of captured adults peaked in late June (Figure 1a). Between 20 and 27 June, large numbers of moths were detected by light traps in Bayannur City, Ordos City, and Ulanqab City in Inner Mongolia, Zhangjiakou City in Hebei, Xinzhou City in Shanxi, and Yanqing District in Beijing. The ovarian developmental stages of moths varied significantly across these sites. Specifically, a total of 44,000 and 33,000 moths were detected by high-light traps in Dalad Banner, Inner Mongolia, and Kangbao County, Hebei, respectively, from 23 to 26 June. On the peak day, a single light trap captured 24,000 and 22,000 moths at these two locations, respectively. On 25 June, a single trap in Chahar Right Front Banner recorded 35,000 *L. sticticalis* moths, with 100% of the dissected females exhibiting ovarian development at 3–4 stages, indicating an immigrant population. In conventional light trap monitoring, the catches on the peak day in late June were 7382, 4656, and 1718 moths in Kangbao County, Wanquan District of Hebei, and Linhe District of Inner Mongolia, respectively.

### 3.2. Migratory Trajectories of Loxostege sticticalis in Northern China

Forward trajectory analysis and probability estimation revealed that, after emigration from 12 monitoring sites—including Daixian (Shanxi), Weichang, Zhuolu, Kangbao, and Wanquan (Hebei), Xinghe, Fengzhen, Dalad Banner, Liangcheng, Urad Front Banner, and Urad Middle Banner (Inner Mongolia), and Jianping (Liaoning)—the potential landing areas of *Loxostege sticticalis* were primarily concentrated in Inner Mongolia, Hebei, and Shanxi, followed by Northeast China and Mongolia (Figure 2a). The migratory patterns of *L. sticticalis* in northern China can be categorized into three main types. Type (1) is southwest–northeast migration from the North China Plain towards Northeast China, with Daixian and Zhuolu representing typical departure areas. For example, moths that took off from Daixian (Shanxi) during the peak stage showed a 63.33% probability of moving through Hebei and Inner Mongolia by the second night. By the fourth and fifth nights, the probabilities of reaching Liaoning and Jilin were 17.49% and 13.81%, respectively, with the farthest eastward dispersal extending to Russia. Similarly, populations from Zhuolu (Hebei) had a 9.55% probability of reaching Liaoning on the fourth and fifth nights. Type (2) is northwestward migration from Liangcheng (Inner Mongolia), Kangbao (Hebei), and Jianping (Liaoning). Taking Kangbao as an example, populations emigrating from this area showed probabilities of entering Mongolia at 16.67%, 32.05%, and 38.16% on the third, fourth, and fifth nights, respectively. Notably, moths from Weichang and Wanquan (Hebei), as well as Xinghe and Fengzhen (Inner Mongolia), exhibited characteristics of both Type (1) and Type (2) migration patterns. For example, those from Weichang had a 2.95% chance of reaching the northernmost point of Heilongjiang after five nights of flying, and moths from all four sites had a chance of moving across the border into Mongolia, with the highest chance being 65.97% (Wanquan). Type (3) is short-distance migration affected by the Yin Mountains barrier, with populations from Dalad Banner, Urad Front Banner, and Urad Middle Banner (Inner Mongolia) mostly moving shorter distances and landing mostly in the autonomous region and along the China–Mongolia border. For example, moths from Urad Middle Banner had a 5.00% probability of landing in Mongolia by the second night.

**Figure 2.**
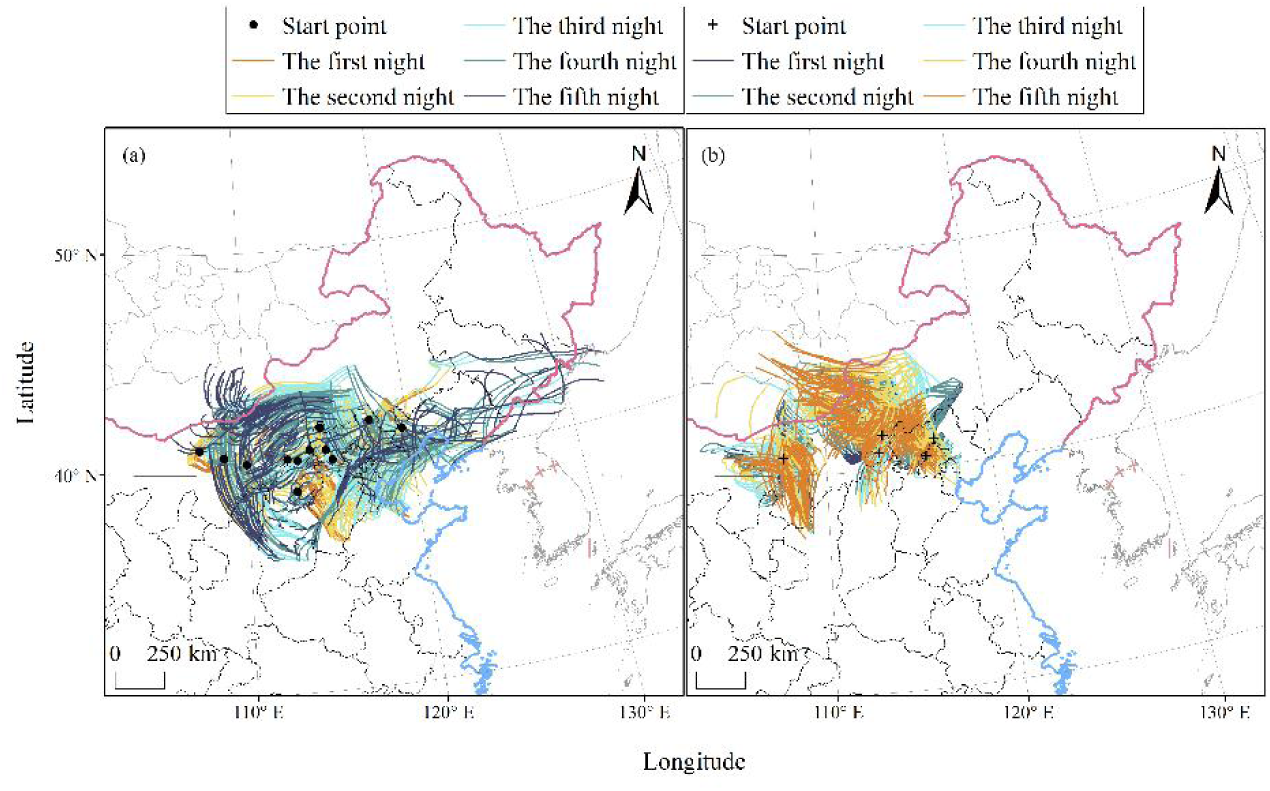
Forward (**a**) and backward (**b**) trajectories of *L. sticticalis* adults in northern China in 2022.

Backward trajectory analysis and probability estimation revealed that, except for the immigrant population in Shangdu (Inner Mongolia) primarily originating from Shanxi, other monitoring sites—including Chahar Right Front Banner and Linhe (Inner Mongolia), Yanqing (Beijing), and Fengning (Hebei)—were influenced to varying degrees by immigrations from Mongolia (Figure 2b). Among these, Chahar Right Front Banner was the primary landing zone for moths from Mongolia, with an overall probability from Mongolia of 18.26%. This was followed by Yanqing, which had a 2.69–6.02% probability of encountering pests from Mongolia over the second to fifth nights. The proportion of immigrant sources from Inner Mongolia in Yanqing was relatively low, higher only than that in Fengning, which was most significantly affected by moths originating from Beijing and southern Hebei, with these regions collectively accounting for 91.13–95.93% of the immigrant populations over the second to fifth nights.

### 3.3. Migratory Pathways and Main Landing Areas of Loxostege sticticalis from Mongolian Source Regions

Simulations of the migratory trajectories of *Loxostege sticticalis* in northern China indicate frequent insect source exchanges between China and Mongolia. To further validate this conclusion, we analyzed the migratory pathways and main landing areas of Mongolian *L. sticticalis* populations based on field survey data from 50 places across 11 provinces in Mongolia: Selenge (1 site), Khentii (4), Dornod (3), Uvurkhangai (3), Bayankhongor (2), Govi-Altai (3), Bulgan (1), Tuv (23), Umnugovi (6), Dornogovi (2), and Dundgovi (2) (Figure 1). These sites cover the perennial occurrence areas of *L. sticticalis* and had an elevation range from 700 to 2200 m above sea level. We also analyzed the migratory pathways and main landing areas of this pest originating from Mongolia. Monitoring data showed that *L. sticticalis* was mainly observed from early June to early August (Figure 1b). Adults were observed at 45 of the surveyed sites (86.54%), and the average moth count per hundred steps at these positive sites was 19. The highest density was recorded in Bayannuur, Bulgan Province (47.86° N; 104.90° E), with 70 moths per hundred steps on 14 June, followed by Altanbulag Soum, Tuv Province (47.88° N; 105.85° E), with 52 moths per hundred steps on the same date.

Further trajectory simulation and probability estimation revealed relatively frequent insect source exchanges between Mongolian populations and those in Russia and China. Between 28 May and 13 August, the Mongolian cross-border migratory populations had a 76.53% chance of entering China, compared to only 23.47% entering Russia. In May and August, the probabilities of *L. sticticalis* migrating into China were as high as 91.35% and 99.35%, respectively. Under suitable wind conditions in June and July, the probabilities of this pest crossing into China notably exceeded 65.06% and 55.60%, respectively. These data indicate a substantially higher probability of Mongolian *L. sticticalis* populations migrating into China. As a critical passageway for all three migratory routes, Inner Mongolia recorded a landing probability of 92.60%, identified as the primary landing area, supplemented by Hebei, Gansu, Shaanxi, Shanxi, and Heilongjiang as affected regions. In May and July, moths advanced into Northeast China passing through Inner Mongolia, exhibiting a distinct “cyclonic” migration pattern. In May, moths passing through Inner Mongolia had a 63.34% probability of reaching Hebei within 2–5 nights, with the farthest extending to Liaoning (Figure 3a). In July, the overall trajectories and farthest landing points were located further north geographically compared to May, as far as Heilongjiang (Figure 3c). In June, the migratory paths from Mongolia were predominantly southeasterly, followed by easterly and southwesterly directions (Figure 3b). The southeasterly routes passed through Inner Mongolia, Shaanxi, Shanxi, and Hebei, with the farthest extending to Shandong. Notably, in late June, the emigration trajectories from Mongolian populations clearly corresponded to the immigration trajectories in northern China (Figure 3e). For instance, moths emigrating from Bayanchandmani on June 24 landed in Chahar Right Front Banner, Inner Mongolia, where a substantial immigrant population was captured on the same day, indicating frequent population exchange between China and Mongolia. The easterly routes in June were similar to the overall July trajectories, extending as far east as Heilongjiang with a landing probability of 3.91%. The southwesterly routes, after passing through Inner Mongolia, could reach Gansu as early as the second night (17.82%), with the farthest trajectories reaching Xinjiang (3.29%). In August, the insects originating from Mongolia only exchanged with those from Inner Mongolia, China (Figure 3d).

**Figure 3.**
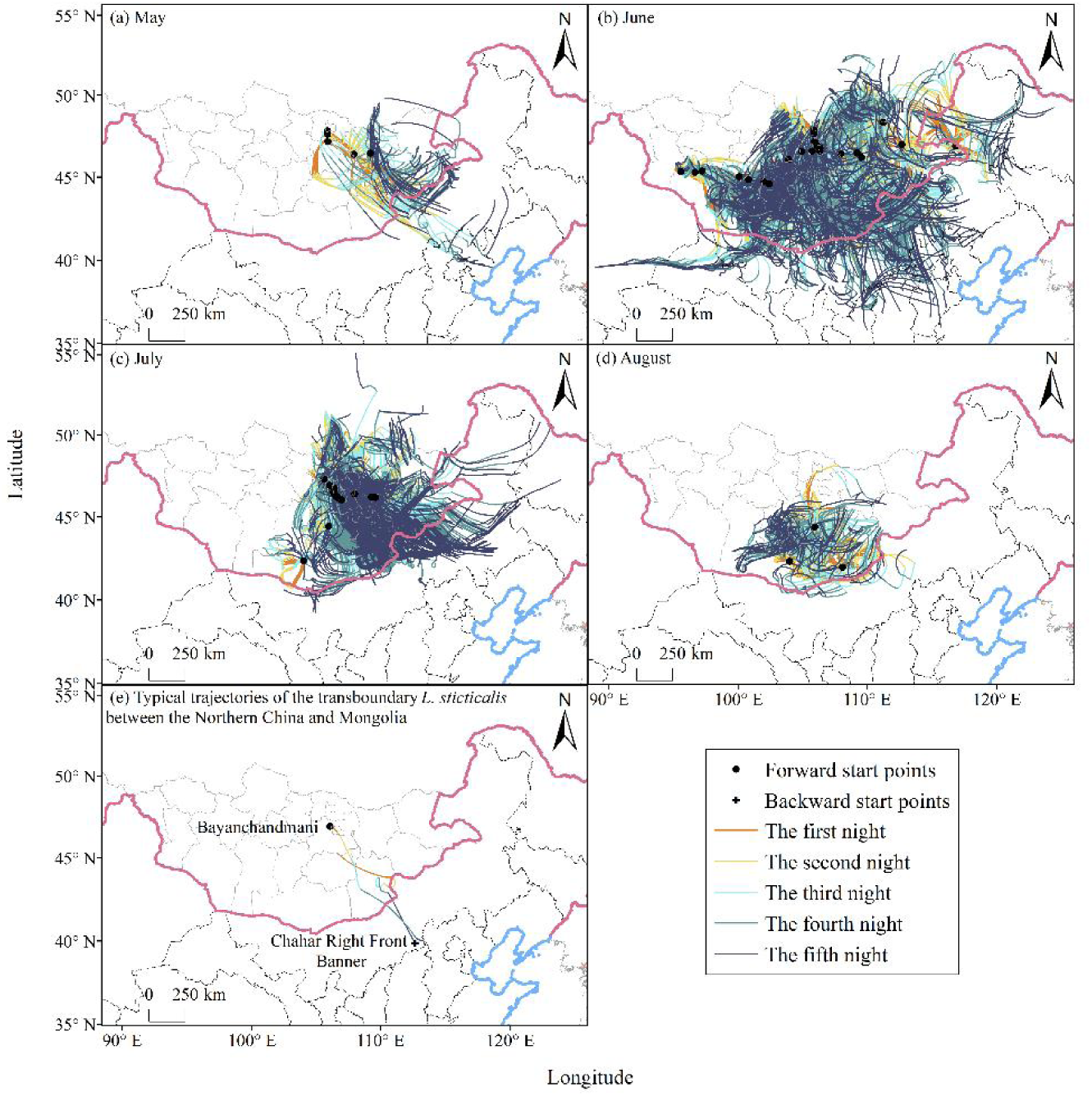
Forward trajectories of migratory *L.sticticalis* in Mongolia in 2022.

Based on this analysis, the migratory pathways of *Loxostege sticticalis* populations from Mongolia into China can be broadly categorized into three types: the first is a southeasterly, quasi-linear path originating from central and eastern Mongolia, crossing the China–Mongolia border, through Inner Mongolia, towards Shanxi and Hebei provinces, extending as far as Shandong; the second is an easterly, “cyclonic” path, where populations from eastern Mongolia move eastward into Inner Mongolia and Northeast China, with the farthest reach extending to Heilongjiang; the third is a southwesterly path, where moths from western Mongolia follow a trajectory detouring around the Altai Mountains, passing through Inner Mongolia and Gansu on the northern side of the Qilian Mountains, with the farthest recorded dispersal reaching Xinjiang.

### 3.4. Synoptic Conditions During the Peak Stages of Loxostege sticticalis Adults in China

#### 3.4.1. Synoptic Situation During the Peak Migration Period of *Loxostege sticticalis*

During 18–20 June 2022, a cold air mass developed on the west of Lake Baikal and had not yet moved southward (Figure 4A). Accompanied by a shortwave trough disturbance, a Mongolian Cyclone formed on 19 June along the western side of the Greater Khingan Mountains and the China–Mongolia–Russia border. From 20–21 June, the center of the cold air mass moved southward, but low-level atmospheric conditions remained dominated by the Mongolian Cyclone, with precipitation areas overlapping perfectly with its center (Figure 4B). This synoptic situation significantly altered the forward trajectories from Weichang and Xinghe (Figure 2a) and restricted the migration of Mongolian insects towards Shangdu County (Figure 2b). By 22 June, an NCCV had developed, with its center located between 40° N and 50° N. From 22 to 24 June, the cold vortex intensified continuously, with its low-pressure center moving southward and extending eastward, while the associated cold center lagged behind, squeezing and eventually absorbing the Mongolian Cyclone by 23 June (Figure 4A). The rainfall zone of marked nocturnal cumulative precipitation expanded on 22 and 23 June, with the main precipitation area covering most of Shandong, eastern Inner Mongolia, central Jilin, southeastern Hebei, and western Liaoning, where nocturnal cumulative precipitation exceeded 30 mm (Figure 4B). By 25 June, the vortex center weakened to 550 hPa, and no considerable widespread precipitation was observed in China. The cold vortex profoundly shaped forward trajectories from Zhuolu, Wanquan, Fengzhen, and Daixian (Figure 2a) and backward trajectories from Chahar Right Front Banner (Figure 2b), making them exhibit an elongated, easterly “cyclonic” pattern. The trajectories during the third to fifth nights of flight were notably longer than those during the first night of migration. This situation was influenced by the Yin Mountains and the eastward movement and weakening of the cold vortex, which led to wind speeds below 5 m/s in the rear sector of the vortex (Figure 4B). Except for the peak intensity of the vortex on 22–23 June, regions in the south of North China and central Inner Mongolia were generally influenced by southerly winds (Figure 4B). This flow pattern facilitated the migration of insects originating from southern areas like Shanxi and Shaanxi towards Linhe, while Fengning and Yanqing were primarily affected by pests from regions like Tianjin (Figure 2b). From 25 to 27 June, the influence of the NCCV gradually weakened. A long trough west of Lake Baikal split into shortwave troughs that followed the weakening cold vortex (Figure 4A). This caused considerable shifts in trajectory directions around 25 June for sites like Kangbao and Liangcheng. After takeoff, moths initially migrated eastward under the cold vortex’s influence, but then shifted northward or even northwestward due to the impact of the shortwave troughs (Figure 2a). After the shortwave trough caught up with the cold vortex on 26 June, precipitation increased by at least 5 mm in North China and central Inner Mongolia (Figure 4B). The squeezing effect of the trough accelerated the movement of the residual cold center after 26 June, displacing it by approximately 5° latitude compared to 24 June and by more than 2° compared to 25 June (Figure 4A). This acceleration substantially increased the migration distance of *L. sticticalis*. However, populations from Dalad Banner, Urad Front Banner, and Urad Middle Banner, obstructed by the topography of the Yin Mountains, primarily engaged in short-distance migration and caused localized damage (Figure 2a).

**Figure 4.**
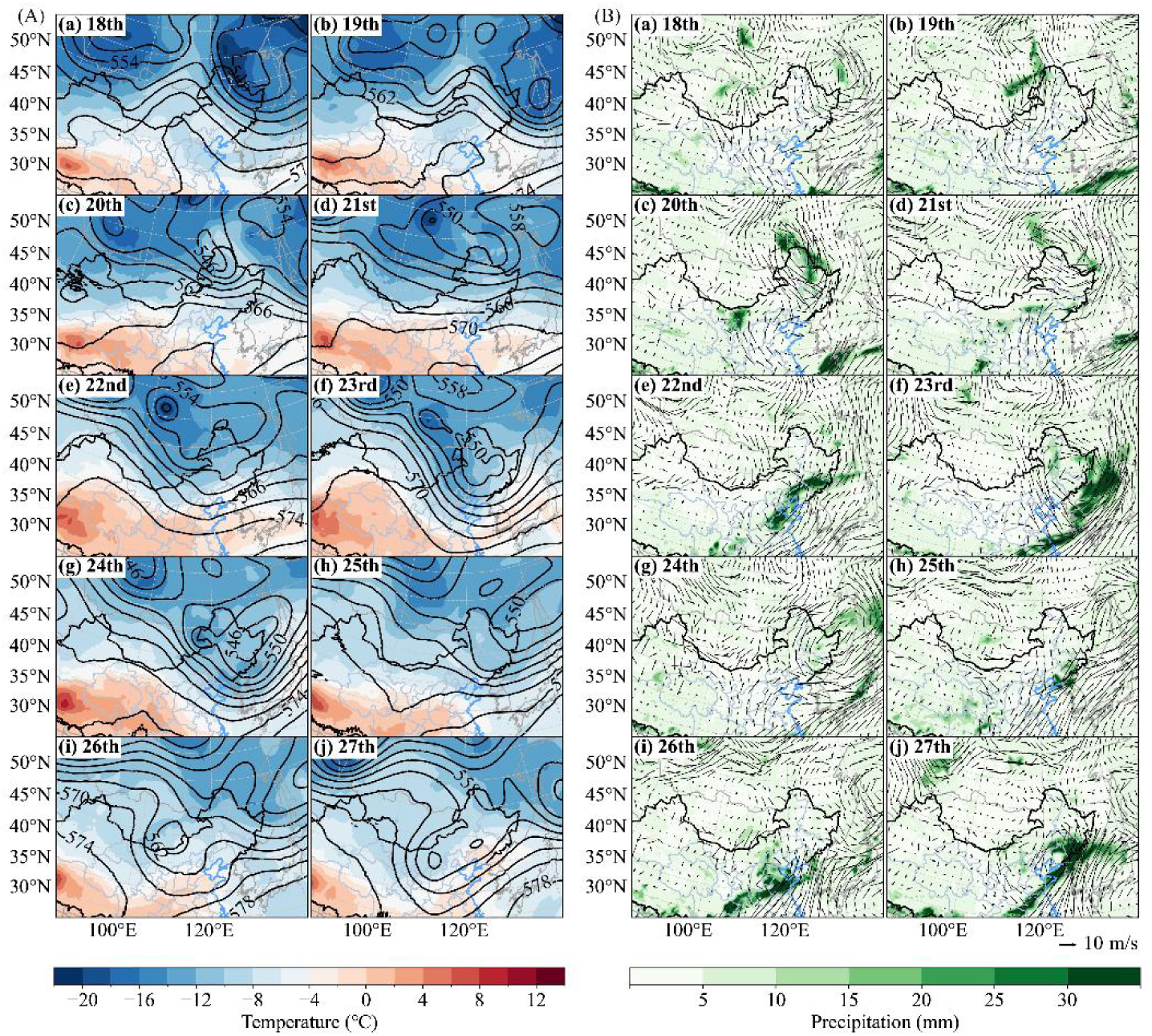
Average nighttime 500 hPa temperature and geopotential height (**A**), and 850 hPa wind field with accumulated precipitation (**B**) during the migration peak of *L. sticticalis* adults from northern China in June 2022.

In summary, aside from topographical influences, the migratory trajectories of *Loxostege sticticalis* populations at most monitoring sites closely corresponded to the development of the Mongolian Cyclone and the NCCV. Moreover, the peak stages of light-trap catches in northern China during this study were primarily driven by the Northeast China Cold Vortex system.

#### 3.4.2. Specific Meteorological Mechanisms Affecting the Migration and Landing of *Loxostege sticticalis*

By analyzing the synoptic conditions on peak days and extracting relevant meteorological elements (including wind speed, wind direction, temperature, downdrafts, and precipitation), the meteorological mechanisms contributing to the large-scale, concentrated landing of *Loxostege sticticalis* in northern China in late June 2022 were further clarified. As shown in Table 1, the average nocturnal wind speed at 850 hPa during the peak adult occurrence period in northern China was only 3.70 ± 2.52 m/s, with a mean wind direction of 168.54 °, indicating prevailing weak southerly winds across the monitoring sites. However, from 22 to 23 June, wind speeds increased significantly in some areas. Zhuolu (Hebei) and Jianping (Liaoning) had mean nighttime wind speeds of 10.27 m/s and 9.66 m/s, respectively. This period coincided with the vigorous phase of the NCCV, which made it easier for *L. sticticalis* to migrate vast distances north and wreak havoc. Additionally, the average nocturnal temperature at 850 hPa across the monitoring sites in late June was 22.19 ± 4.28 °C, above the low temperature threshold for *L. sticticalis* flight. This indicates that temperature did not restrict migratory activity during this period. On nights without precipitation, downdrafts were observed at 77.78% of the sites. The maximum vertical velocity occurred in Zhuolu (Hebei) on the 23rd, reaching 28.05 × 10^−2^ Pa/s. Regarding precipitation, Weichang and Fengning (Hebei) experienced moderate rain and drizzle, respectively, on the nights of 20–21 June. Between 26 and 27 June, four sites—Liangcheng (Hebei), Wanquan (Hebei), Yanqing (Beijing), and Dalad Banner (Inner Mongolia)—experienced rainfall events, mainly moderate rain. In summary, downdrafts and precipitation were identified as the primary meteorological causes for the concentrated landing of *L. sticticalis* in northern China.

**Table 1.** Average nighttime meteorological factors at stations on peak days of *L. sticticalis* captured by light traps.

| Monitoring Location | Date | 850 hPa Wind Speed/(m/s) | 850 hPa Wind Direction/° | 850 hPa Temperature/°C | Accumulated Precipitation/m | 850hPa Vertical Velocity/(×102 Pa/s) | Precipitation Class | Category of Searchlight Traps |
| --- | --- | --- | --- | --- | --- | --- | --- | --- |
| Shangdu County | 22 June 2022 | 2.75 | 114.13 (SE) | 17.74 | 0.00 | 2.68 |  | Surface |
| Urad Middle Banner | 23 June 2022 | 5.76 | 177.99 (S) | 24.36 | 0.00 | −16.64 |  | Aerology |
| Dalad Banner | 23 June 2022 | 5.77 | 106.08 (E) | 24.18 | 0.00 | 23.68 |  | Aerology |
| Dalad Banner | 26 June 2022 | 2.25 | 171.22 (S) | 17.94 | 1.15 | 4.39 | Light rain | Surface |
| Linhe District | 25 June 2022 | 4.47 | 154.97 (SE) | 28.63 | 0.00 | −22.71 |  | Surface |
| Chahar Right Front Banner | 25 June 2022 | 1.33 | 216.82 (SW) | 25.24 | 0.00 | 3.18 |  | Aerology |
| Xinghe County | 25 June 2022 | 1.77 | 199.40 (S) | 25.58 | 0.00 | 3.81 |  | Aerology, Surface |
| Fengzhen City | 25 June 2022 | 1.40 | 231.37 (SW) | 25.00 | 0.00 | 1.23 | Drizzle | Aerology |
| Liangcheng County | 25 June 2022 | 1.16 | 286.76 (W) | 27.96 | 0.00 | 2.07 | Drizzle | Surface |
| Liangcheng County | 26 June 2022 | 1.11 | 60.75 (NE) | 17.30 | 5.93 | −1.38 | Moderate rain | Aerology |
| Fengning County | 20 June 2022 | 5.01 | 221.43 (SW) | 22.10 | 0.05 | −5.87 | Drizzle | Surface |
| Zhuolu County | 23 June 2022 | 10.27 | 125.18 (SE) | 20.38 | 0.00 | 28.05 |  | Surface |
| Dai County | 24 June 2022 | 2.21 | 267.56 (W) | 26.82 | 0.00 | 8.09 |  | Aerology |
| Dai County | 25 June 2022 | 1.22 | 294.93 (NW) | 27.98 | 0.00 | 10.59 | Drizzle | Surface |
| Weichang County | 21 June 2022 | 2.12 | 240.02 (SW) | 16.59 | 11.77 | −1.04 | Moderate rain | Aerology |
| Kangbao County | 25 June 2022 | 3.29 | 20.30 (N) | 20.86 | 0.00 | 5.79 |  | Aerology, Surface |
| Wanquan District | 26 June 2022 | 3.69 | 208.10 (SW) | 17.45 | 11.74 | −11.63 | Moderate rain | Aerology, Surface |
| Yanqing District | 27 June 2022 | 5.16 | 21.67 (N) | 16.44 | 0.47 | −20.62 | Light rain | Aerology |
| Jianping County | 25 June 2022 | 9.66 | 83.64 (E) | 18.64 | 0.73 | 2.19 | Light rain | Aerology |

### 3.5. Interactive Effects of the NCCV and Mongolian Cyclone During the Peak Stages of Loxostege sticticalis

In northern China and Mongolia, the NCCV and the Mongolian Cyclone interact and influence each other. A cold trough prior to the formation of the cold vortex often triggers Mongolian Cyclones. These processes collectively disrupt the synoptic background, thereby influencing the migration and landing of *Loxostege sticticalis*. This study systematically examined the occurrence processes of the NCCV and the Mongolian Cyclone, along with their interactive effects, to further investigate the large-scale atmospheric circulation patterns governing the extensive migration and collective landing of *L. sticticalis* from 28 May to 13 August 2022. During this period, there were seven NCCV events and three Mongolian Cyclone events. Among the NCCV processes, three were classified as Central Vortex (40–50° N) and four as Northern Vortex (50–60° N), with no Southern Vortex (35–40° N) processes observed (Table 2). Analysis revealed that all three Mongolian Cyclones were either influenced or triggered by shortwave troughs from the cold center preceding the formation of the NCCV.

**Table 2.**
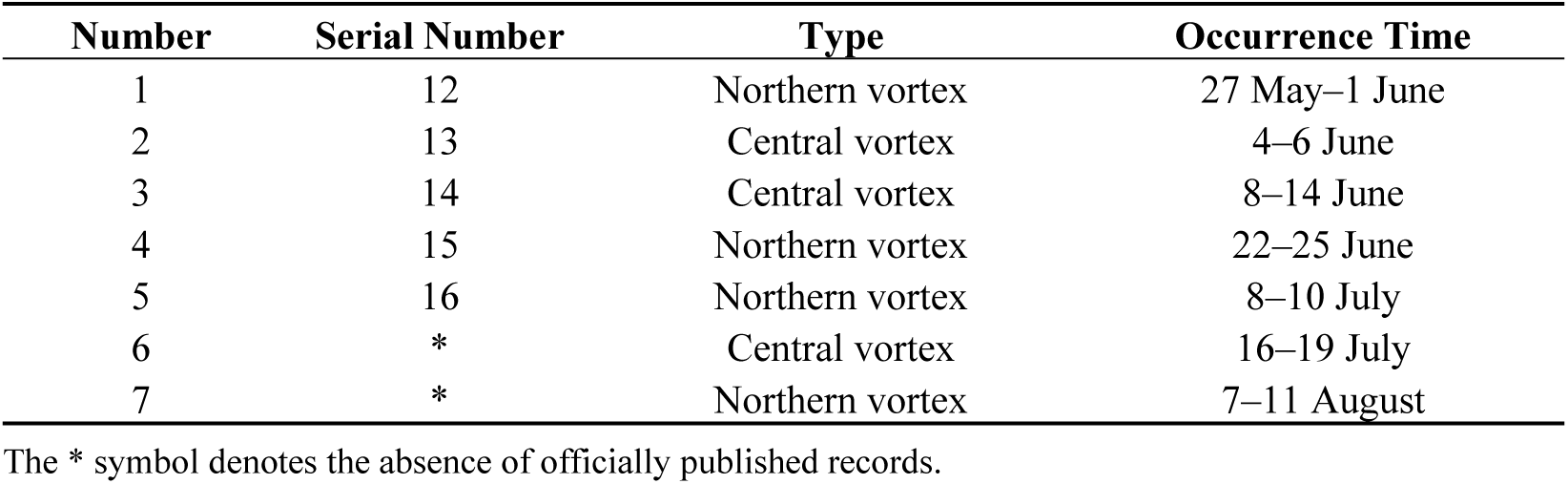
Occurrence of the NCCV from May to August 2022. The serial numbers are cited from the Liaoning Meteorological Service (http://ln.cma.gov.cn/xwzx/ztbd/dblwjcybgb/index.html accessed on 5 December 2024). The NCCV is classified into southern (35–40° N), central (40–50° N), and northern types (50–60 °N) [24].

| Number | Serial Number | Type | Occurrence Time |
| --- | --- | --- | --- |
| 1 | 12 | Northern vortex | 27 May–1 June |
| 2 | 13 | Central vortex | 4–6 June |
| 3 | 14 | Central vortex | 8–14 June |
| 4 | 15 | Northern vortex | 22–25 June |
| 5 | 16 | Northern vortex | 8–10 July |
| 6 | * | Central vortex | 16–19 July |
| 7 | * | Northern vortex | 7–11 August |

The \* symbol denotes the absence of officially published records.

From 28 to 31 May, the migratory trajectories, influenced by Cold Vortex No.12 in central and eastern Mongolia, exhibited a “cyclonic” pattern (Figure 5a). Transported downwind to the Greater Khingan Mountains, their further migration was hindered by northerly winds (Figure 3a). Three typical nocturnal synoptic processes during May-August 2022 were identified, occurring on 22 June, 12 July, and 4 August. On 22 June, the NCCV and a Mongolian Cyclone coexisted, with temperature fields at both upper and lower levels coupled while geopotential height fields were inconsistent. The night of the 21st was dominated by the Mongolian Cyclone (Figure 6a,b), and then the NCCV was dominant on 22–23 June. Cyclonic activity was most pronounced in June, with two Mongolian Cyclones and three NCCV processes recorded, including three Central Vortex and one Northern Vortex process. The frequent alternation of these two systems led to a mean weak wind field which, in turn, induced the greatest variety of immigrant pathways and the highest volume of *L. sticticalis* immigration. Throughout June, the 850 hPa level along the China–Mongolia border was consistently dominated by northerly–northwesterly winds at night, while southerly winds in North China were less obstructive than those in other months (Figure 5b), which provided favorable transport conditions for the migration of Mongolian *L. sticticalis* into China. Precipitation in the border zone totaled less than 25 mm for the month, while that in northern Shanxi, Hebei, and Beijing exceeded 25 mm, and southern Hebei and Shandong saw over 75 mm of precipitation (Figure 5b). Consequently, part of the populations in central and eastern Mongolia migrated southeastward through Inner Mongolia, reaching as far as Shandong, while another part was carried into Northeast China by the “cyclonic” wind field (Figure 3b). Populations from western Mongolia, entering China north of the Qilian Mountains in Gansu, moved from east to west towards Xinjiang under weak wind shear, experiencing less than 5 mm of precipitation during this period (Figure 5b). In early to mid-July, the Central Vortex, Northern Vortex, and Mongolian Cyclones occurred in succession. Northern Vortex No. 16 was particularly notable for its extensive coverage, great depth, and high intensity. Although this vortex began to weaken since 11–12 July, its residual influence, combined with the Mongolian Cyclone it helped spark, strongly dominated during early to mid-July (Figure 6c,d). During this period, populations from central and eastern Mongolia were carried into Northeast China by the “cyclonic” wind field (Figure 3c). No cold vortices or Mongolian Cyclones developed in late July. Marked wind shear and strengthened wind fields were observed along the border in July, with prevailing southerlies over northern China and northerlies over Mongolia. This pattern not only enhanced cross-border insect source exchange but also facilitated the concentrated landing of airborne *L. sticticalis* (Figure 5c). Concurrently, monthly cumulative precipitation exceeded 25 mm in Shanxi, Hebei, the three northeastern provinces, and eastern Inner Mongolia (Figure 5c), providing favorable conditions for moth landing. In the first half of August, one Northern Vortex process was recorded, but no Mongolian Cyclones developed. On 4 August, a transient, shallow mesoscale cyclone formed, triggered by the cold trough prior to cold vortex genesis (Figure 6e,f), which was soon replaced by the cold vortex that developed on 7 August. Consequently, wind directions were highly variable in early August, leading to shorter and more concentrated migration distances, but in diverse directions, exhibiting a “cyclonic” characteristic (Figure 3d). During this period, sustained northerly–northwesterly winds prevailed over Mongolia along the border, while adjacent areas in China experienced southerly winds (Figure 5d), together leading to massive landings of Mongolian *L. sticticalis* along the border region.

**Figure 5.**
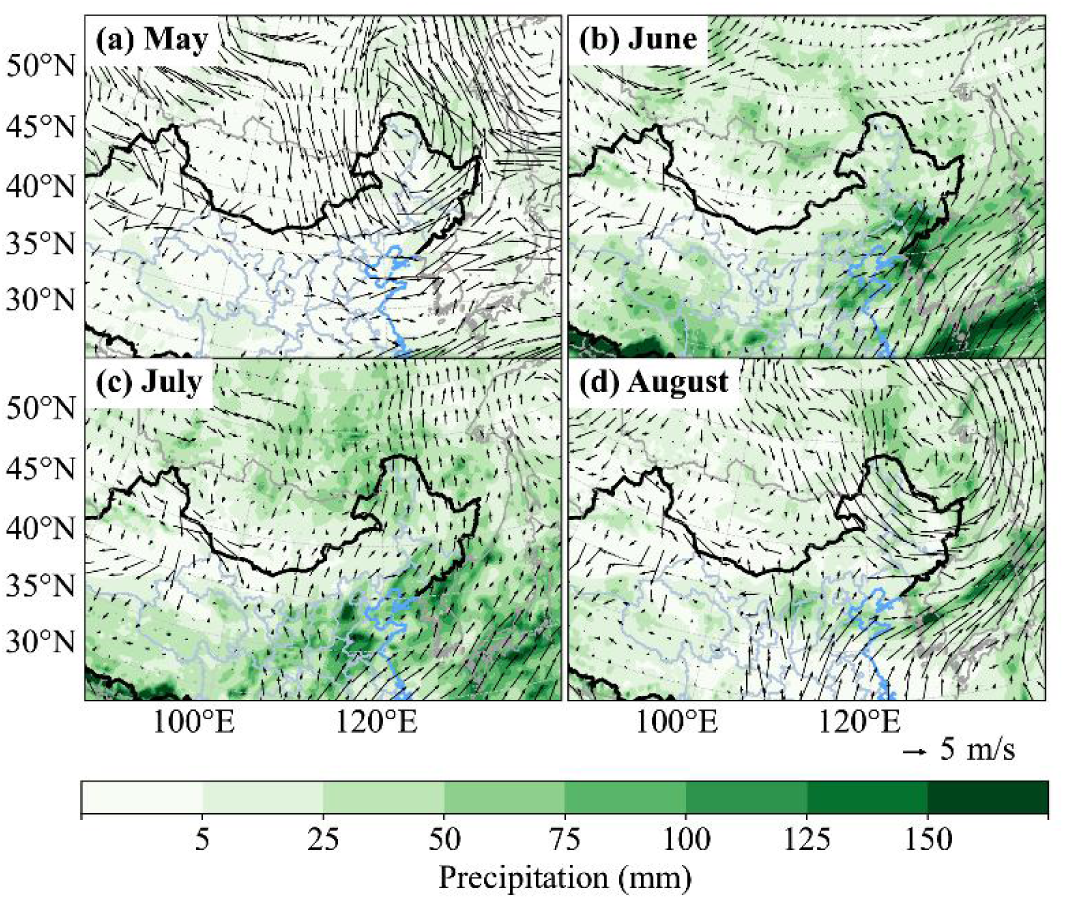
Average nighttime wind field at 850 hPa and accumulated precipitation during the migration peak of *L. sticticalis* from May to August in 2022.

**Figure 6.**
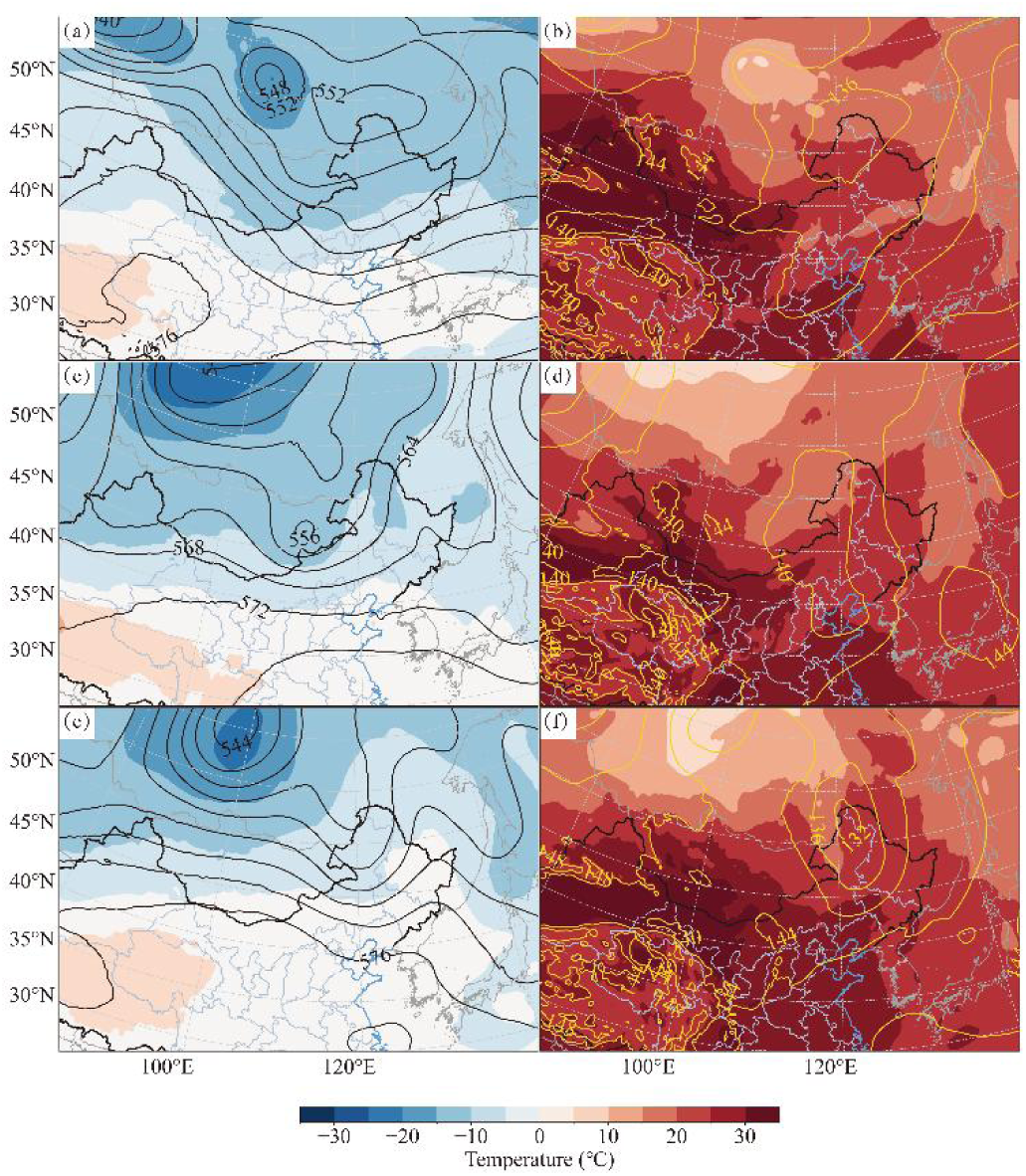
Average nighttime temperature and geopotential height at 500 hPa (**a**,**c**,**e**) and 850 hPa (**b**,**d**,**f**) during the typical NCCV occurrence during May to August in 2022. a and b are before the cold vortex absorbed the Mongolian Cyclone on 22 June; (**c**,**d**) are the Mongolian Cyclone spawned by the trough behind the NCCV on 12 July; (**e**,**f**) are troughs before the cold vortex, giving rise to a cyclone on 4 August.

Throughout the period from May to August 2022, as shown by the mean nocturnal wind field at 850 hPa (Figure 5), the peak flight period of adults was characterized by predominant northerly winds over Mongolia and southerly winds over northern China. The persistent wind shear along the China–Mongolia border, in terms of its spatiotemporal distribution and intensity, played a decisive role in shaping the migratory pathways, distance, and landing zones of *L. sticticalis*. The NCCV and Mongolian Cyclone systems, which gave rise to and controlled this large-scale atmospheric circulation pattern, along with their interactions, were identified as the main reasons for this pest migrating from Mongolia to northern China during the study period.

## 4. Discussion

### 4.1. Notable Exchange of Insect Populations Between China and Mongolia

Since the beginning of the 21st century, it has been widely recognized that, in addition to local overwintering populations, insects colonizing Northeast China are partially derived from Mongolia and Russia [6,16,17,19]. In 2015, Frolov classified *L. sticticalis* into two geo-population systems, the European–Western Siberian (Russia, Ukraine, and Kazakhstan) and the Central Asian (China and Mongolia) populations, based on geographical features of the Cayan and Altai Mountains. He proposed the East Siberian corridor as a considerable topographic zone facilitating population connectivity between China and Mongolia [20]. Data from a field survey conducted in 2022 identified that primary infestation areas were mainly located in the border areas of Inner Mongolia, northern Hebei, and northern Shanxi in China, as well as in central and eastern Mongolia (Figure 1). This spatial distribution is closely related to the region’s ecological conditions. *L. sticticalis* is mostly found in China north of the 12 °C mean annual temperature isotherm and in areas with yearly precipitation of 100–700 mm. The worst damage happens in the 2nd–3rd generation occurrence zones, which have a mean annual temperature between 0 and 8 °C, making them favorable for the pest to overwinter [46,47]. This area lies in the temperate dry and semi-arid zone of China [48], with most of the land covered in grasslands and desert steppes, and a higher coverage of forestland cropland in the eastern region [48,49]. Likewise, the infestations of *L. sticticalis* in Mongolia are found in temperate dry or semi-arid temperature zones. Most of Mongolia’s terrain is covered in grasslands and desert steppes, but there are also some forest steppes in the north [50,51], which are generally less disturbed by humans. In 2022, the average January temperatures of all monitoring sites in Mongolia were above the average supercooling point for diapausing *L. sticticalis* larvae (−26.79 °C) [52]. The mean annual temperature range was between −1.21 °C and 7.58 °C, allowing development for 1–3 generations of the pest per year [46]. The average annual precipitation across the sites was 278.72 mm, with 86.05% of the sites meeting the annual precipitation criteria (100–700 mm) for suitable *L. sticticalis* habitats. During the 2022 migration window, 54.55% of the sites met the precipitation requirement for medium-to-high suitability during the emergence period (15–50 mm) (Table S2) [18]. In conclusion, the core outbreak areas in both China and Mongolia share high consistency in climate types, underlying surface composition, and key meteorological factors, together forming core infestation areas for *L. sticticalis* in both countries.

Based on survey data from the two core areas, this study conducted trajectory simulations. The forward trajectory results showed that populations from North China (e.g., Daixian in Shanxi and Zhuolu in Hebei) primarily migrated northeastward towards Northeast China, while populations from North China (exemplified by Kangbao in Hebei) and Inner Mongolia also migrated north–northwestward, towards the China–Mongolia border. Backward trajectory analysis indicated that the principal sources of *L. sticticalis* populations in northern China included Hebei, Inner Mongolia, and Mongolia. At the same time, the potential emigration trajectories of Mongolian populations showed that *L. sticticalis* from Mongolia may go directly southeast into China after departure if the weather is good. In addition, the Northeast Cold Vortex and the Mongolian Cyclone could induce Mongolian populations to take another two migratory routes: one towards the east and one towards the southwest. These routes extended as far south as northwestern Shandong, north to northern Heilongjiang, and west to southern Xinjiang. Notably, in late June 2022, the emigration paths from Mongolia clearly aligned with the backward trajectories of populations in northern China (Figure 3e). Our findings are consistent with the inferences drawn from the backward trajectories in North China by Chen et al. [6]. Chen et al. [6,16] pointed out that a large-scale immigration of *L. sticticalis* occurred in Northeast China in late May and early June 2001, which was earlier than the occurrence of populations in North China, suggesting that these adults are highly likely to have come from northeastern Mongolia. Furthermore, in 2002 and 2007, the insects in Northeast China mainly originated from eastern Mongolia, and their migration paths closely aligned with the identified emigration paths from Mongolia in late May and early June [6]. The migratory path along the border areas of Shanxi, Inner Mongolia, and Hebei in early August 2003 also coincided with the emigration path from southern Mongolia during the period described here [16].

Ovarian development is an important physiological indicator for determining characteristics of migratory insects, applied in species like the armyworm moth (*Mythimna separata*) [53], rice leafroller (*Cnaphalocrocis medinalis*) [54], brown rice planthopper (*Nilaparvata lugens*) [55], and fall armyworm (*Spodoptera frugiperda*) [56]. According to long-term population monitoring data, the sustained presence of only individuals with low ovarian development stages suggests the emigration of the local population. In contrast, a prominent appearance of high-grade individuals among declining local population density typically indicates immigration from external sources. When the proportion of low-grade individuals gradually decreases while the number of high-grade individuals consistently increases, this generally indicates the natural development of a local population [57]. Restricted by limited data in 2022, this study could not undertake long-term monitoring or systematic ovarian dissection at all of the monitoring sites, and thereby failed to thoroughly characterize the population dynamics of *Loxostege sticticalis*. Future research should concentrate on enhancing the systematic surveillance of *L. sticticalis* populations and further examining its occurrence patterns and small-to-medium-scale transboundary migration routes. This will provide a theoretical foundation for improving the prediction, forecasting, and scientific control of *L. sticticalis* in China.

### 4.2. Noteworthy Impact of the NCCV

Meteorological analysis further revealed that the overall low-level wind fields in June and July were highly conducive to the invasion of *Loxostege sticticalis* from Mongolian source regions and the insect source exchange between Mongolia and Russia. Notably, during June, when the NCCV and the Mongolian Cyclone frequently interacted, an additional southwesterly migration route was generated, originating from western Mongolia, extending as far as southern Xinjiang in China. Existing surveys indicate that Xinjiang is a high-altitude occurrence area for *L. sticticalis* in China, with insects there primarily affecting Hotan in southern Xinjiang, as well as areas bordering Kazakhstan including Altay, Bole, and Tacheng in northern Xinjiang [43,58]. In southern Xinjiang, the peak abundance period of overwintering-generation adults typically occurs in early July [59]. Therefore, substantial numbers of *L. sticticalis* migrating from Mongolia into Xinjiang via this southwesterly route (“Mongolia–Xinjiang, China”) in June supplement the local population base and potentially exacerbate outbreak threats in some areas. Moreover, studies of the average nighttime wind patterns during the peak days in late May and early to mid-August showed that cyclonic systems can change the background wind patterns for *L. sticticalis* migration, making it easier for insects to move between central and eastern Mongolia and China. Previous studies have confirmed that both the Mongolian Cyclone and the NCCV play major roles in the migration process of *L. sticticalis*. The low-level wind direction associated with these systems influences migration direction, while the wind speed determines migration distance. Precipitation and strong convective weather triggered by these cyclonic systems can also lead to massive landings of *L. sticticalis* [15,16,60]. In this study, we found that from June to August 2022, the NCCV relatively dominated the synoptic patterns. This is because the NCCV develops deeply over Northeast China, Inner Mongolia, and Mongolia, exerting substantial and prolonged effects on local wind fields [24,32,33]. The changes in weather conditions and meteorological elements caused by the NCCV’s interaction with and influence over the Mongolian Cyclone substantially affected the migration pathways, distances, and the occurrence of landings of *L. sticticalis* in that year. Moreover, by leveraging these specific weather conditions, *L. sticticalis* from Russia could also migrate into China and cause damage. Given suitable wind fields and the presence of substantial source populations in Russia and Mongolia, China faces an ongoing risk of successive immigration and potential outbreaks of *L. sticticalis* [12,19,61]. In conclusion, our findings underscore the regulatory role of mid- to high-latitude cyclonic systems in East Asia on the cross-border migration of *L. sticticalis* between China and Mongolia.

### 4.3. Effects of Climate Change and Extreme Rainfall Events on the Movement of Loxostege sticticalis

Temperature and precipitation have a significant effect on how *Loxostege sticticalis* behaves. The best temperature for laying eggs is between 19 and 25 degrees Celsius, with 22 degrees Celsius being optimal for migration [30]. Adults will not fly if the temperature is below 15 °C, but they tend to fly significantly more on a large scale if the temperature is over 20 °C [8,31]. Global warming may lead to an earlier emergence of *L. sticticalis* [47]. The regions that experience the most substantial warming, which are the main infestation areas of *L. sticticalis* in China, have seen an annual temperature increase 1.5 to 3 times that of other areas [62]. Monitoring data from the Pest Forecasting Division of the National Agro-Tech Extension and Service Center confirmed that, in 2022, the first appearance of overwintering adults under light traps occurred earlier than that in average years in most parts of China [63]. Two to three generations can be completed every year in China, with adults appearing successively at monitoring sites across various provinces from late April to mid-May [46]. The overwintering generation peaks in mid- to late June [64], while the first generation’s peak has progressively delayed in recent years, now occurring from mid- to late June to early August [14]. In 2022, the peak stages of the overwintering generation occurred in mid- to late June, slightly later than that in average years. In Mongolia during the same year, temperatures at each monitoring site reached the developmental threshold of 10.5 °C for *L. sticticalis* [46] between the end of April and mid- to late May [46]. The population fluctuation trends observed at Mongolian monitoring sites in 2022 were similar to those in northern China, with peak periods occurring slightly earlier than those in China, and the number of moths flushed per 100 paces in the field showing two peaks: one in early to mid-June and the other in early to mid-July. In 2022, the infestation was less severe in China than in 2021 [65]. One probable factor for this is comparatively higher precipitation in predominant infestation areas in that year. Global warming is intensifying rainfall (high confidence), and for every 1 °C rise in temperature, extreme daily precipitation events are anticipated to climb by about 7% (high confidence) [62]. Consistent low temperatures and wet weather can further decrease female moths’ ability to reproduce and enhance their sterility [47]. For instance, the end of the second and third outbreak cycles of *L. sticticalis* in China coincided with periods of low temperatures, heavy rainfall, and severe adverse weather conditions [60]. However, the impact of climate warming on precipitation exhibits considerable regional variation [66]. Although precipitation intensity increased in northeastern and northern China in 2022 [66], the frequency of local droughts also rose in some areas, such as northwestern China [67], thereby affecting the habitat suitability for *L. sticticalis* populations. High temperature and humidity during summer can inhibit the early development of eggs and larvae [47]. For instance, in provinces like Shandong and Hunan, such conditions may explain the rare reports of major outbreaks [68,69]. Nevertheless, against the backdrop of continued climate warming, the narrowing temperature discrepancy between northern and southern China and reduced precipitation in some southern areas could favor the occurrence, damage, and further northward migration of *L. sticticalis*, provided that the pests successfully pass their early developmental stages.

### 4.4. Relationship Between Agricultural Landscape Changes and Loxostege sticticalis Outbreaks

Some researchers, based on the cyclical patterns of *Loxostege sticticalis* outbreaks and field population monitoring data, along with solar activity cycles, have inferred that the fourth outbreak cycle of *L. sticticalis* occurred approximately between 2016 and 2025 [11,70]. Although the population levels remained low in 2016 and 2017 [9,71], a resurgence was observed between 2018 and 2019 [10,11], culminating in a typical severe outbreak year in 2020 [12], marking the entry into the fourth outbreak cycle. *L. sticticalis* has mostly stayed at moderate levels in recent years, without major outbreaks, thanks to timely monitoring, early warnings, and good control. Areas including Inner Mongolia, Hebei, and the three northeastern provinces where major food crops are cultivated suffered severe infestations [72]. Hebei and Shanxi, located in the climatically suitable Yellow River basin, feature vast areas of staple crops like corn and millet. In the majority of the 96 banners and counties in Inner Mongolia, primary host crops of *L. sticticalis*, including wheat, corn, potatoes, and soybeans, are grown. The principal cultivation regions of soybeans and potatoes correspond with the primary occurrence zones of *L. sticticalis* identified in this study [73]. Mongolia shares similar land cover types with northern China, with its agricultural structure primarily focused on livestock farming, although wheat (a staple food crop), potatoes, and other crops are also grown [18]. Adjustments in the planting structure of host plants have influenced the migration and occurrence patterns of *L. sticticalis* during its fourth cycle [11,70]. *L. sticticalis* is polyphagous but exhibits distinct feeding and oviposition preferences. Its preference for soybeans and potatoes is significantly higher than for corn. During oviposition, adults lay significantly more eggs on leguminous forages than on gramineous forages [74]. Survey data from the China Agricultural and Rural Information Network (http://www.agri.cn/ accessed on 6 April 2025) indicate that Heilongjiang and Inner Mongolia were major soybean-producing provinces in 2022. Heilongjiang, Jilin, Inner Mongolia, Henan, Liaoning, Shanxi, Gansu, and the Ningxia Hui Autonomous Region are key corn-growing areas. Soybean production has increased steadily year by year since 2015, while corn production remained roughly the same. The distribution of crops significantly influences infestation degrees and areas. Since agricultural structure adjustments, the environment has become more conducive to the migration, survival, and reproduction of *L. sticticalis*. Therefore, sustained and close monitoring of *L. sticticalis* population dynamics in China holds considerable importance for guiding early warning and sustainable management of this major pest in the context of evolving agricultural landscapes.

## 5. Conclusions

In 2022, the principal infestation areas of *Loxostege sticticalis* were located in the border regions of Inner Mongolia, northern Hebei, and northern Shanxi in China, in addition to central and eastern Mongolia, with frequent exchanges between populations in China and Mongolia. The migrating populations from Mongolia followed three main pathways: a predominant southeasterly route, with supplementary eastward “cyclonic” and southwesterly paths. The main landing areas in China were North China and Northeast China, with migration range extending as far as Shandong, Heilongjiang, and Xinjiang. Populations from North China (such as Daixian in Shanxi and Zhuolu in Hebei) primarily migrated northeastward toward Northeast China, while populations from North China (exemplified by Kangbao in Hebei) and Inner Mongolia also migrated north–northwestward, heading toward the China–Mongolia border. Insect sources from North China could reach Northeast China and Mongolia after 1–5 consecutive nights of migration. Meteorological systems played a major regulatory role in the cross-border migration of *L. sticticalis*. From June to August 2022, there were seven NCCV events and three Mongolian Cyclone events. During the migration period, the interactions between these systems determined the synoptic circulation. Their spatial and temporal distribution and intensity were very important at the very stage when the pathways were established and the range of the cross-border migration was expanded. Persistent wind shear at the China–Mongolia border made it easier for insects to interact between the two areas. Wind fields of prevailing NCCV events provided the driving force for the immigration of Mongolian populations into China and for the northeastward migration of populations from North China, contributing to the risk of infestation. Moreover, the precipitation and downdrafts triggered by the NCCV were key factors driving the concentrated landing of *L. sticticalis* populations in northern China.

## Supplementary Materials

**Table S1:**
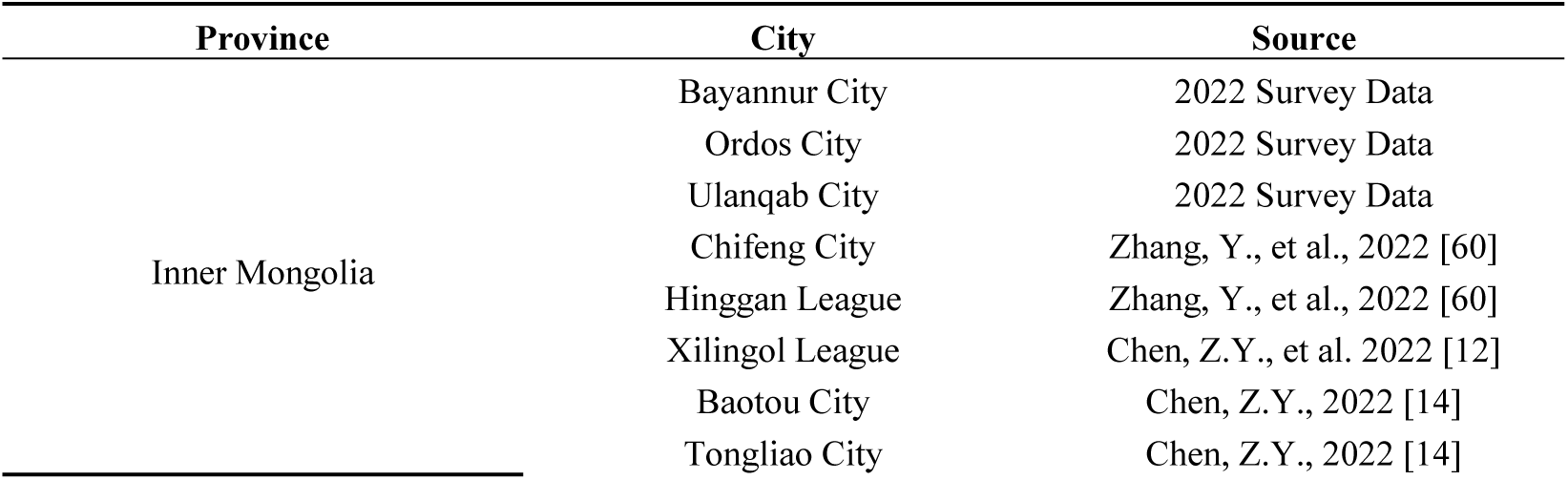
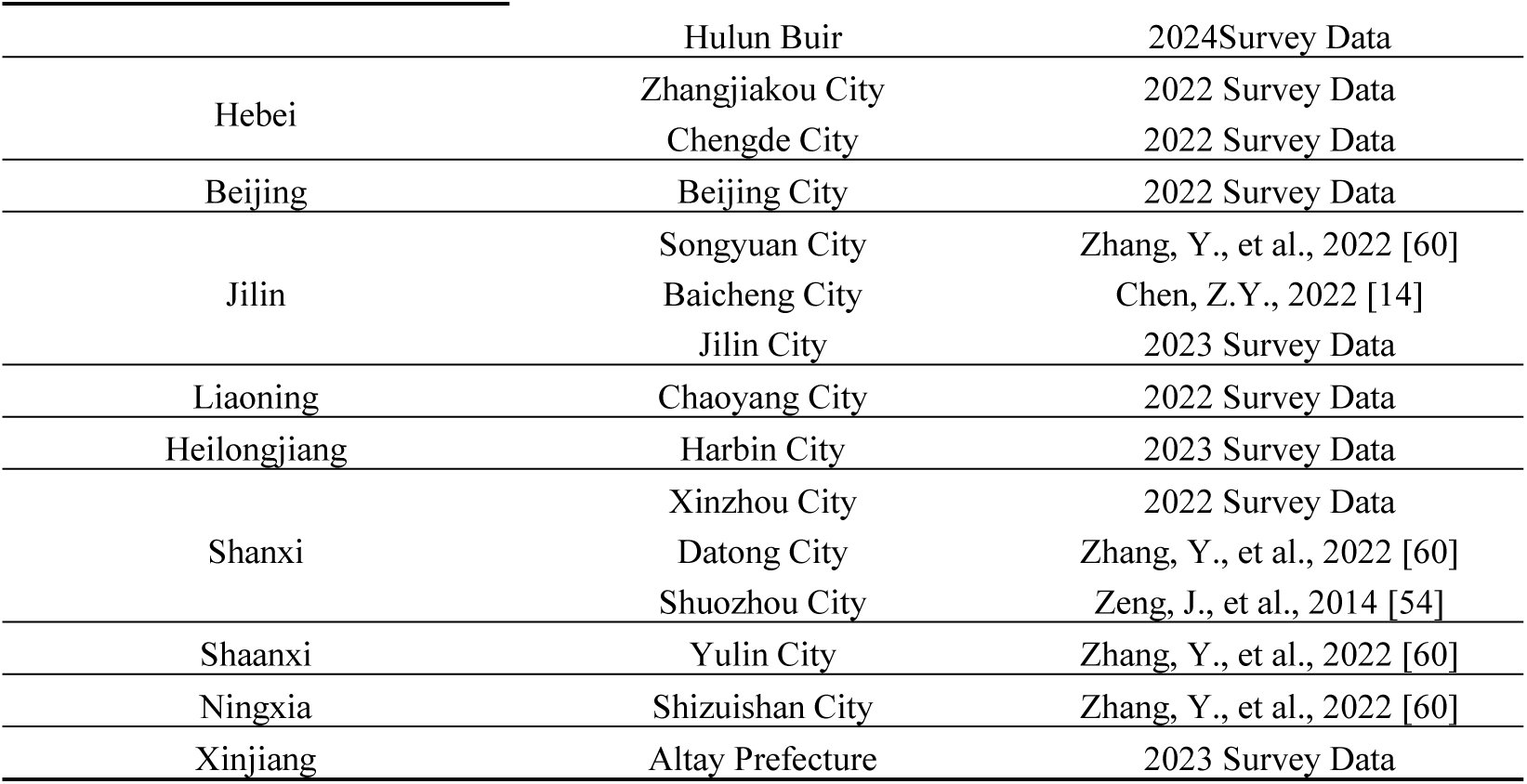
Valid insect source areas of *L. sticticalis* in northern China in recent years.

| Province | City | Source |
| --- | --- | --- |
| Inner Mongolia | Bayannur City | 2022 Survey Data |
|  | Ordos City | 2022 Survey Data |
|  | Ulanqab City | 2022 Survey Data |
|  | Chifeng City | Zhang, Y., et al., 2022 [60] |
|  | Hinggan League | Zhang, Y., et al., 2022 [60] |
|  | Xilingol League | Chen, Z.Y., et al. 2022 [12] |
|  | Baotou City | Chen, Z.Y., 2022 [14] |
|  | Tongliao City | Chen, Z.Y., 2022 [14] |
|  | Hulun Buir | 2024Survey Data |
| Hebei | Zhangjiakou City | 2022 Survey Data |
|  | Chengde City | 2022 Survey Data |
| Beijing | Beijing City | 2022 Survey Data |
| Jilin | Songyuan City | Zhang, Y., et al., 2022 [60] |
|  | Baicheng City | Chen, Z.Y., 2022 [14] |
|  | Jilin City | 2023 Survey Data |
| Liaoning | Chaoyang City | 2022 Survey Data |
| Heilongjiang | Harbin City | 2023 Survey Data |
| Shanxi | Xinzhou City | 2022 Survey Data |
|  | Datong City | Zhang, Y., et al., 2022 [60] |
|  | Shuozhou City | Zeng, J., et al., 2014 [54] |
| Shaanxi | Yulin City | Zhang, Y., et al., 2022 [60] |
| Ningxia | Shizuishan City | Zhang, Y., et al., 2022 [60] |
| Xinjiang | Altay Prefecture | 2023 Survey Data |

**Table S2:**
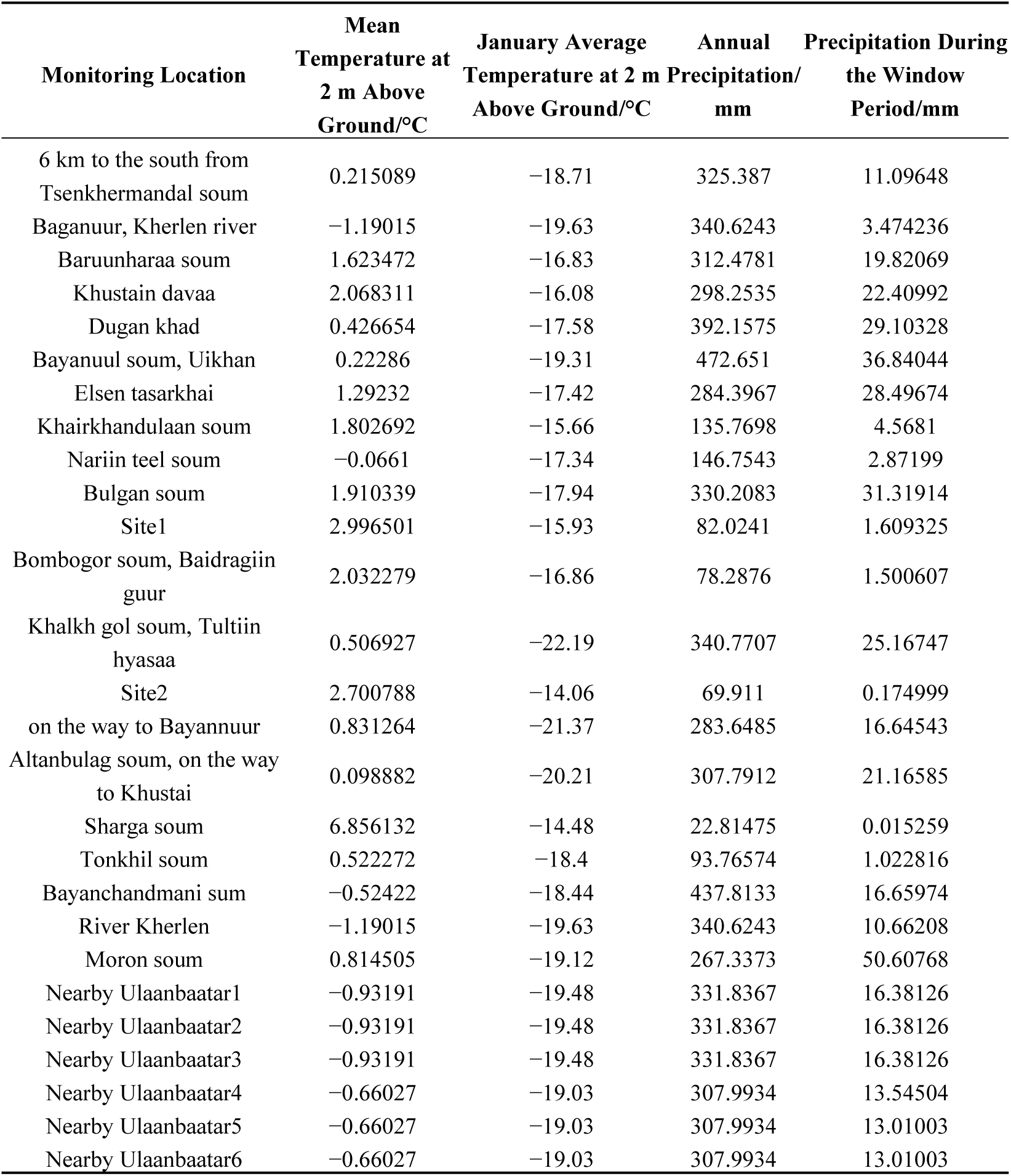
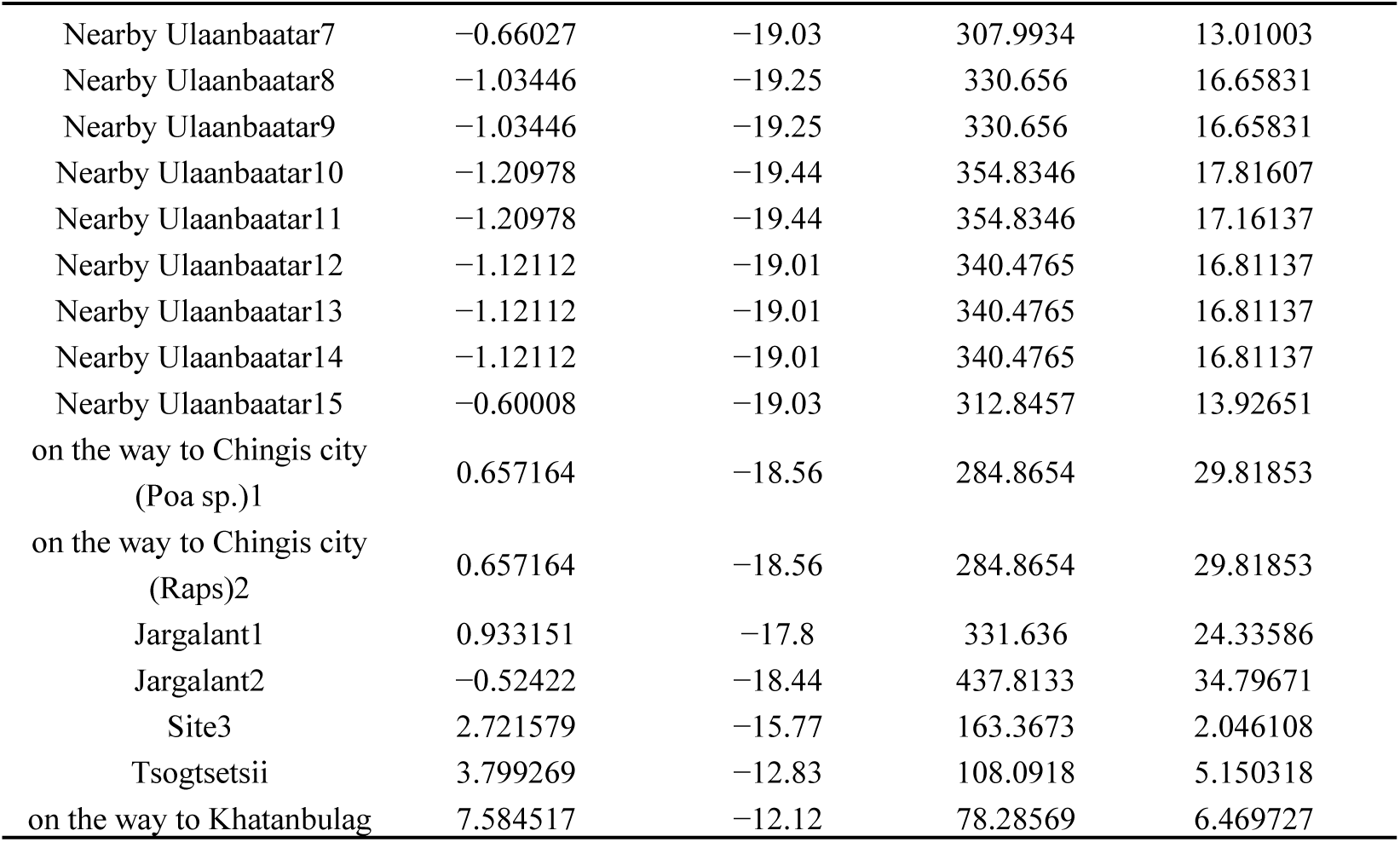
Habitat suitability in the core areas of Mongolia in 2022.

## Author Contributions

Conceptualization, Q.-L.W., F.-J.C., and G.-J.W.; methodology, Q.-L.W. and X.-Y.P.; formal analysis, X.-Y.P.; investigation, Y.-Y.Z.; resources, Q.-L.W.; writing—original draft preparation, X.-Y.P., R.Z., and H.-B.G.; writing—review and editing, Q.-L.W., G.-J.W., and F.-J.C.; visualization, X.-Y.P.; supervision, Q.-L.W.; funding acquisition, Q.-L.W., G.-J.W., and F.-J.C. All authors have read and agreed to the published version of the manuscript.

## Data Availability Statement

The meteorological data supporting the results of this study are publicly available on the open-access website https://cds.climate.copernicus.eu/ [accessed on 16 December 2024], as mentioned in the Materials and Methods section. The insect data are presented in Figure 1 of the article. For further inquiries regarding the data or additional information, please contact the corresponding author.

## Abbreviations

The following abbreviations are used in this manuscript:

L. sticticalis: Loxostege sticticalis
NCCV: Northeast China Cold Vortex
HYSPLIT: Hybrid Single-Particle Lagrangian Integrated Trajectory
AGL: Above ground level

## Disclaimer/Publisher’s Note

The statements, opinions and data contained in all publications are solely those of the individual author(s) and contributor(s) and not of MDPI and/or the editor(s). MDPI and/or the editor(s) disclaim responsibility for any injury to people or property resulting from any ideas, methods, instructions or products referred to in the content.

## References

1. National Scientific Research Cooperation Group of Loxostege sticticalis. Research on occurrence, forecast and control of *Loxostege sticticalis*. China Plant Prot. 1987, S1, 1–9. (In Chinese)

2. Afonin, A.N.; Akhanaev, Y.B.; Frolov, A.N. The range of the beet webworm *Loxostege sticticalis* L. (Lepidoptera, Pyraloidea: Crambidae) in the former USSR territory and its subdivision by the number of generations per season. Entomol. Rev. 2014, 94, 200–204.

3. Petrukha, O.I.; Tribel’ S.A. The population dynamic of *Loxostege sticticalis*. Zashchita Rastenii 1975, 4, 41–43.

4. Tian, S.Y. Research on control of *Loxostege sticticalis*. Acta Agric. Boreali-Sin. 1963, 3, 15–22. (In Chinese)

5. Luo, L.Z.; Li, G.B.; Cao, Y.Z. The 3rd occurrence cycle of the beet webworm *Loxostege sticticalis* may be coming in China. Plant Prot. 1996, 22, 50–51. (In Chinese)

6. Chen, X.; Zhai, B.P.; Gong, R.J.; Yin, M.H.; Zhang, Y.; Zhao, K.J. The source area of spring populations of meadow moth, Loxostege sticticalis L. (Lepidoptera: Pyralidae) in northeastern China. Acta Ecol. Sin. 2008, 28, 1521–1535. (In Chinese)

7. Ministry of Agriculture and Rural Affairs of the People’s Republic of China. Circular of the Ministry of Agriculture and Rural Affairs and Ministry of Finance on Announcing the List of Advantageous Characteristic Industrial Clusters to be Constructed in 2020. 2020. Available online: https://www.gov.cn/zhengce/zhengceku/2020-05/22/content_5513870.htm (accessed on 20 September 2023).

8. Luo, L.Z.; Cheng, Y.X.; Tang, J.H.; Zhang, L.; Jiang, X.F. Temperature and relative humidity are the key factors for population dynamics and outbreak of the beet webworm, Loxostege sticticalis. Plant Prot. 2016, 42, 1–8. (In Chinese)

9. Zeng, J.; Jiang, Y.Y.; Liu, J. The regional pattern of *Loxostege sticticalis* L. varied during a new occurrence intermission in China. Acta Ecol. Sin. 2018, 38, 1832–1840. (In Chinese)

10. Liu, J.; Jiang, Y.Y.; Zeng, J.; Chen, Y.; Wang, C.R.; Zhang, Y.H.; Tao, Y.L. Meadow moth *Loxostege sticticalis* occurred severely in partial area of northeast of China in 2018. China Plant Prot. 2019, 39, 36–41. (In Chinese)

11. Jiang, X.F.; Zhang, L.; Cheng, Y.X.; Jiang, Y.Y.; Liu, J. The fourth occurrence cycle of the beet webworm *Loxostege sticticalis* may be coming in China. Plant Prot. 2019, 45, 79–81. (In Chinese)

12. Chen, Z.Y.; Zhang, Z.; Liu, J.; Kang, A.G.; Zhao, S.M.; Yin, X.J.; Li, Z.Q.; Xie, A.T.; Zhang, Y.H. Spatiotemporal dynamics and source of *Loxostege sticticalis* in Northern China in 2020. Sci. Agric. Sin. 2022, 55, 907–919. (In Chinese)

13. Feng, H. By 2020, the planting area will be expanded to more than 100 million mu and 30% of potatoes will be used as staple food. Rural. Areas Agric. Farmers 2016, 3, 19. (In Chinese)

14. Chen, Z.Y. Analysis of Migration and Meteorological Background for Meadow Moths (Loxostege sticticalis L.) in North and Northest China. Master’s Thesis, Chinese Academy of Agricultural Sciences, Beijing, China, 2022. (In Chinese)

15. Sun, H.Y. Relationship of northeast cold vortex with migration and landing of *Loxostege sticticalis* (Ls). J. Meteorol. Environ. 2014, 30, 85–89. (In Chinese)

16. Chen, X. The Migration, Overwinter Regulation and Outbreak Mechanism of Meadow Moth (*Loxostege sticticalis* L.). Ph.D. Thesis, Nanjing Agricultural University, Nanjing, China, 2010. (In Chinese)

17. Huang, S.Z. Study on Spatiotemporal Population Dynamics of *Loxostege sticticalis* (Lepidoptera: Pyralidae) in China. Ph.D. Thesis, Chinese Academy of Agricultural Sciences, Beijing, China, 2010. (In Chinese)

18. Frolov, A.N.; Malysh, Y.M.; Tokarev, Y.S. Biological features and population density forecasts of the beet webworm *Pyrausta sticticalis* L. (Lepidoptera, Pyraustidae) in the period of low population density of the pest in Krasnodar Territory. Entomol. Rev. 2008, 88, 666–675.

19. Luo, L.Z.; Huang, S.Z.; Jiang, X.F.; Zhang, L. Characteristics and causes for the outbreaks of beet webworm, *Loxostege sticticalis* in northern China during 2008. Plant Prot. 2009, 35, 27–33. (In Chinese)

20. Frolov, A.N. The beet webworm *Loxostege sticticalis* L. (Lepidoptera, Crambidae) in the focus of agricultural entomology objectives: I. The periodicity of pest outbreaks. Entomol. Rev. 2015, 95, 147–156.

21. Bell, J.R.; Aralimarad, P.; Lim, K.S.; Chapman, J.W. Predicting insect migration density and speed in the daytime convective boundary layer. PLoS ONE 2013, 8, e54202.

22. Wainwright, C.E.; Reynolds, D.R.; Reynolds, A.M. Linking small-scale flight manoeuvers and density profiles to the vertical movement of insects in the nocturnal stable boundary layer. Sci. Rep. 2020, 10, 1019.

23. Nieto, R.; Gimeno, L.; De la Torre, L.; Ribera, P.; Gallego, D.; García-Herrera, R.; García, J.A.; Nuñez, M.; Redaño, A.; Lorente, J. Climatological features of cutoff low systems in the Northern Hemisphere. J. Clim. 2005, 18, 3085–3103.

24. Sun, L.; Zheng, X.Y.; Wang, Q. The climatological characteristics of Northeast cold vortex in China. *Q*. J. Appl. Meteorol. 1994, 5, 297–303. (In Chinese)

25. Chen, X. Trajectory Analysis of Immigration Population of Meadow Moth in Northeast Part of China. Master’s Thesis, Northeast Agricultural University, Harbin, China, 2003. (In Chinese)

26. Luo, L.P.; Xue, M.; Zhu, K.F. The initiation and organization of a severe hail-producing mesoscale convective system in east China: A numerical study. J. Geophys. Res. Atmos. 2020, 125, e2020JD032606.

27. Yin, L.; Ping, F.; Xu, H.B.; Chen, B.J. Numerical simulation and the underlying mechanism of a severe hail-producing convective system in east China. J. Geophys. Res. Atmos. 2021, 126, e2019JD032285.

28. Hu, P.Y.; Chen, C.L.; Lin, H.F.; Xu, S.; Kong, L.J.; Liu, Q.; Ji, Y.M.; Zhao, Z.Q. Definition and characteristics analysis of the intensity of Northeast Cold Vortex. J. Meteorol. Environ. 2021, 37, 100–105. (In Chinese)

29. Liu, W. Climatic characters of Northeast China Cold Vortex and its impact on precipitation in Inner Mongolia. Meteorol. J. Inn. Mong. 2021, 2, 9–13. (In Chinese)

30. Tang, J.H.; Cheng, Y.X.; Sappington, T.W.; Jiang, X.F.; Zhang, L.; Luo, L.Z. Egg hatch and survival and development of beet webworm (Lepidoptera: Crambidae) larvae at different combinations of temperature and relative humidity. J. Econ. Entomol. 2016, 109, 1603–1611.

31. Sun, Y.J.; Gao, Y.B. Discussion on beet webworm sources of migration and spring occurrence. In Proceedings of the 60th Anniversary and Symposium of Entomological Society of China, Chongqing, China, 4 October 2004. (In Chinese)

32. Xie, Z.W.; Bueh, C. Cold vortex events over northeast China associated with the yakutsk-okhotsk blocking. Int. J. Climatol. 2017, 37, 381–398.

33. Liu, C.H.; Fang, Y.H.; Zhang, K.; Li, Y.N.; Wang, Y. Northeast China cold vortex is the key factor influencing the high-impact agroclimatic events in northeast China. *Dyn*. Atmos. Ocean. 2024, 107, 101477.

34. Huang, X.; Bueh, C.; Xie, Z.W.; Gong, Y.F. Mongolian cyclones that influence the northern part of China in spring and their associated low-frequency background circulations. *Chin*. J. Atmos. Sci. 2016, 40, 489–503. (In Chinese)

35. Zhang, Y.H.; Chen, L.; Cheng, D.F.; Jiang, Y.Y.; Lv, Y. The migratory behaviour and population source of the first generation of the meadow moth, *Loxostege sticticalis* L. (Lepidoptera: Pyralidae) in 2007. Acta Entomol. Sin. 2008, 51, 720–727. (In Chinese)

36. Chen, Z.Y.; Zhang, Z.; Zhang, Y.H. Progress in research on monitoring and forecasting the occurrence of the beet webworm, Loxostege sticticalis. Chin. J. Appl. Entomol. 2021, 58, 552–564. (In Chinese)

37. Zhang, Y.; Wang, L.J.; Li, X.R.; Zhang, A.H.; Zhang, Y.H. Simulation of *Loxostege sticticalis* migratory flight paths with three trajectory models. *Chin*. J. Appl. Entomol. 2023, 60, 903–912. (In Chinese)

38. Zhu, M.; Radcliffe, E.B.; Ragsdale, D.W.; MacRae, I.V.; Seeley, M.W. Low-level jet streams associated with spring aphid migration and current season spread of potato viruses in the US norther Great Plains. Agric. For. Meteorol. 2006, 138, 192–202.

39. Westbrook, J.K.; Nagoshi, R.N.; Meagher, R.L.; Fleischer, S.J.; Jairam, S. Modeling seasonal migration of fall armyworm moths. Int. J. Biometeorol. 2016, 60, 255–267.

40. Wu, Q.L.; Westbrook, J.K.; Hu, G.; Lu, M.H.; Liu, W.C.; Sword, G.A.; Zhai, B.P. Multiscale analyses on a massive immigration process of *Sogatella furcifera* (Horváth) in south-central China: Influences of synoptic-scale meteorological conditions and topography. Int. J. Biometeorol. 2018, 62, 1389–1406.

41. Wang, X.Y.; Ma, H.T.; Wu, Q.L.; Zhou, Y.; Zhou, L.H.; Xiu, X.Z.; Zhao, Y.C.; Wu, K.M. Comigration and interactions between two species of rice planthopper (*Laodelphax striatellus* and *Sogatella furcifera*) and natural enemies in eastern Asia. Pest Manag. Sci. 2023, 79, 4066–4077.

42. Feng, H.Q.; Wu, K.M.; Cheng, D.F. Spring migration and summer dispersal of *Lorostege sticticalis* (Lepidoptera: Pyralidae) and other insects with radar in northern China. Environ. Entomol. 2004, 33, 1253–1265.

43. Wang, H.; Mu, C.; Ni. Y.F.; Lin, J.; Yu, F.; Wu, L.N.; Xu, G.Q.; Ji, R. Study on *Loxostege sticticalis* ovary development and its resource in Altay Xinjiang. Xinjiang Agric. Sci. 2011, 48, 1324–1328. (In Chinese)

44. Song, W.Y.; Cheng, Y.X.; Luo, L.Z.; Zhang, L.; Wu, J.X.; Jiang, X.F. Effects of migration on reproduction and population outbreak in the beet webworm, Loxostege sticticalis. Plant Prot. 2016, 42, 26–30. (In Chinese)

45. Chen, R.L.; Bao, X.Z.; Wang, S.Y.; Sun, Y.J.; Li, L.Q.; Liu, J.R. An observation on the migratory of, meadow moth by radar. J. Plant Prot. 1992, 19, 171–174. (In Chinese)

46. Luo, L.Z.; Li, G.B. Effective accumulated temperature of meadow moth and division of the generation area. Acta Entomol. Sin. 1993, 36, 332–339. (In Chinese)

47. Tang, J.H. Responses and Adaptation of the Beet Webworm, *Loxostege sticticalis* (Lepidoptera: Carambidiae) to the Variations in Temperature and Humidity. Ph.D. Thesis, Chinese Academy of Agricultural Sciences, Beijing, China, 2016. (In Chinese)

48. Huang, Y.; Liu, X.M.; Yuan, J.F.; Fu, Z.; Qiao, Q. Spatial and temporal changes of carbon storage and its influencing factors in arid and semi-arid region of North China from 2000 to 2020. *Res*. Environ. Sci. 2024, 37, 849–861. (In Chinese)

49. Zhao, A.Z.; Xu, R.H.; Zou, L.D.; Zhu, X.F. Response of grassland vegetation growth to drought in Inner Mongolia of China from 2002 to 2020. Atmosphere 2023, 14, 1613.

50. Chang, S.; Wu, B.F.; Yan, N.N.; Davdai, B.; Nasanbat, E. Suitability assessment of satellite-derived drought indices for Mongolian grassland. Remote Sens. 2017, 9, 650.

51. Vova, O. Assessment of Drought in Grasslands: Spatio-Temporal Analyses of Soil Moisture and Extreme Climate Effects in Southwestern Mongolia. Ph.D. Thesis, Georg-August-Universität Göttingen, Göttingen, Germany, 2021.

52. Li, C.X.; Luo, L.Z.; Pan, X.L. Cold-hardiness in the diapause and non-diapause larvae of meadow moth, Loxostege sticticalis L. Plant Prot. 2006, 2, 41–44. (In Chinese)

53. Luo, L.Z.; Li, G.B.; Hu, Y. Relationship between flight capacity and oviposition of oriental armyworm moths, *mythimna separata* (walker). Acta Entomol. Sin. 1995, 38, 284–289.

54. Zhang, X.X.; Lu, Z.Q.; Geng, J.G.; Li, G.Z.; Chen, X.L.; Wu, X.W. Research on migration pathways of *Cnaphalocrocis medinalis*. Acta Entomol. Sin. 1980, 2, 130–140.

55. Cheng, X.N.; Chen, R.C.; Xi, X.; Yang, L.M.; Zhu, Z.L.; Wu, J.C.; Qian, R.G.; Yang, J.S. Studies on the migrations of brown planthopper *Nilaparvata lugens* Stål. Acta Entomol. Sin. 1979, 22, 1–20.

56. Zhao, S.Y.; Yang, X.M.; He, W.; Zhang, H.W.; Jiang, Y.Y.; Wu, K.M. Ovarian development gradation and reproduction potential prediction in *Spodoptera frugiperda*. Plant Prot. 2019, 45, 28–34.

57. Qi, G.J.; Lu, F.; Hu, G.; Wang, F.Y.; Gao, Y.; Lv, L.H. The application of ovarian dissection in the research on migratory insects in China. China Plant Prot. 2011, 31, 18–22.

58. Li, G.H.; Aerziguli·Aisai; Zhang, L.; Yang, J.F.; Han, Q. Causes and control strategies of meadow moth *Loxostege sticticalis* outbreaks in Xinjiang. Xinjiang Agric. Sci. Technol. 2011, 3, 26. (In Chinese)

59. Zeng, J.; Jiang, Y.Y. Occurrence characteristics and causes of the meadow moth *Loxostege sticticalis* in China in 2012. Plant Prot. 2014, 40, 142–148. (In Chinese)

60. Chen, X.; Jiang, Y.Y.; Meng, Z.P.; Chen, Kuo.; Kang, A.G.; Li, C.M.; Zhai, B.P. Extreme climate has become an important factor causing the termination of outbreak periods of *Loxostege sticticalis* (Lepidoptera: Pyralidae) in China. Acta Entomol. Sin. 2016, 59, 1363–1375. (In Chinese)

61. Huang, S.Z.; Zhang, L.; Xie, D.J.; Tang, J.H.; Jiang, Y.Y.; Mijidsuren, B.; Baasan, M.; Luo, L.Z.; Jiang, X.F. Transboundary migration of *Loxostege sticticalis* (Lepidoptera: Crambidae) among China, Russia and Mongolia. Pest Manag. Sci. 2024, 80, 4650–4664.

62. IPCC. Climate Change 2021: The Physical Science Basis. 2022. Available online: https://www.ipcc.ch/report/ar6/wg1 (accessed on 6 October 2023).

63. National Agro-Tech Extension and Service Center. The Spring Populations of *Loxostege sticticalis* Emerged in a Peak, and One Generation of Larvae Caused Damage in Partial Area. 2022. Available online: https://www.natesc.org.cn/news/des?id=d6df73af-5198-45c3-95eb-0787df3882f6&CategoryId=07e72766-0a38-4dbd-a6a3-c823ce1172bd (accessed on 20 June 2023).

64. Zeng, J.; Jiang, Y.Y. Analysis on occurrence of the meadow moth *Loxostege sticticalis* in China in 2014. Plant Prot. 2016, 42, 194–199. (In Chinese)

65. Zhang, Y.Y.; Liu, J.; Zhao, S.M.; Yin, X.J.; Zhang, Y.H.; Bian, Y.; Zeng, J.; Jiang, Y.Y. Occurrence characteristics and causes of meadow moth *Loxostege sticticalis* outbreaks in China in 2021. *Chin*. J. Appl. Entomol. 2022, 59, 1372–1384. (In Chinese)

66. Shu, Z.K.; Li, W.X.; Zhang, J.Y.; Jin, J.L.; Xue, Q.; Wang, Y.T.; Wang, G.Q. Historical changes and future trends of extreme precipitation and high temperature in China. Strateg. Study CAE 2022, 24, 116–125. (In Chinese)

67. Zhai, P.M.; Zhou, B.Q.; Chen, Y.; Yu, R. Several new understandings in the climate change science. Clim. Change Res. 2021, 17, 629–635. (In Chinese)

68. Wang, W.Y. Poputation Dynamics and Control Strategies of *Loxostege sticticalis* L. (Lepidoptera: Pyralidae) in Lawn in Changsha area. Master’s Thesis, Hunan Agricultural University, Changsha, China, 2006. (In Chinese)

69. Song, H.Y.; Li, L.L.; Zhang, Q.Q.; Sun, C.K.; Li, C.; Lu, Z.B.; Zhu, Z.G.; Yu, Y.; Men, X.Y. Community structure of insects in Shandong province as revealed by searchlight and ground light-traps. *Chin*. J. Appl. Entomol. 2021, 58, 724–735. (In Chinese)

70. Zhang, L.; Jiang, X.F. Occurrence tendency and management strategies of the beet webworm, Loxostege sticticalis in China. Plant Prot. 2022, 48, 68–72. (In Chinese)

71. Jiang, Y.Y.; Liu. W.C.; Huang, C.; Liu, J.; Yang, Q.P.; Lu, M.H. Occurrence trend forecast of major crop diseases and insect pests in China in 2017. China Plant Prot. 2017, 37, 45–49+57. (In Chinese)

72. Zhang, L.; Jiang, X.F. Current state of research on the beet webworm, Loxostege sticticalis (Linnaeus) in China. Chin. J. Appl. Entomol. 2023, 60, 1654–1668. (In Chinese)

73. Wang, F. Research on the regional pattern of staple crops in Inner Mongolia. Inn. Mong. Stat. 2015, 6, 17–18. (In Chinese)

74. Qu, Y.F.; Wang, Q.Q.; Xiao, C.; Li, K.B.; Cao, Y.Z.; Yin, J. Preferences of *Loxostege sticticalis* for oviposition on different pasture plants and the effects of different host plants on the population growth of this pest. *Chin*. J. Appl. Entomol. 2023, 60, 1679–1687. (In Chinese)

